# Lipid-specific protein oligomerization is regulated by two interfaces in Marburg virus matrix protein VP40

**DOI:** 10.1101/2020.11.13.381350

**Authors:** Souad Amiar, Monica L. Husby, Kaveesha J. Wijesinghe, Stephanie Angel, Nisha Bhattarai, Bernard S. Gerstman, Prem P. Chapagain, Sheng Li, Robert V. Stahelin

## Abstract

Marburg virus major matrix protein (mVP40) dimers associate with anionic lipids at the plasma membrane and undergo a dynamic and extensive self-oligomerization into the structural matrix layer which confers the virion shape and stability. Using a myriad of *in vitro* and cellular techniques, we present a mVP40 assembly model highlighting two distinct oligomerization interfaces (N-terminal domain (NTD) and C-terminal domain (CTD)) in mVP40. Cellular studies of NTD and CTD oligomerization interface mutants demonstrated the importance of each interface in the mVP40 matrix assembly through protein trafficking to the plasma membrane and homo-multimerization that induced protein enrichment, plasma membrane fluidity changes and elongations at the plasma membrane. A novel APEX-TEM method was employed to closely assess the ultrastructural localization of and formation of viral particles for wild type and mutants. Taken together, these studies present a mechanistic model of mVP40 oligomerization and assembly at the plasma membrane during virion assembly.

## Introduction

The *Filoviridae* family of viruses, which includes Marburg virus (MARV) and its cousin Ebola virus (EBOV), have been responsible for several highly fatal outbreaks since the late 1960s (Suzuki and Gojobori, 1997; Slenczka and Klenk, 2007; Leroy, Gonzalez and Baize, 2011; Breman *et al.*, 2016; World Health Organization, 2019). Filoviruses are lipid-enveloped viruses harboring a negative sense RNA genome which bud and release new filamentous viral particles from the host cell plasma membrane (Beer, Kurth and Bukreyev, 1999; Kolesnikova *et al.*, 2002; Noda *et al.*, 2002; Kolesnikova, Bamberg, *et al.*, 2004; Bray and Geisbert, 2005; Leroy, Gonzalez and Baize, 2011; Bharat *et al.*, 2012). The viral matrix protein VP40 of MARV and EBOV (mVP40 and eVP40, respectively) is the primary viral component responsible for directing the assembly and budding of viral particles from the host cell plasma membrane inner leaflet (Feldmann, Klenk and Sanchez, 1993; Harty *et al.*, 2000; Jasenosky *et al.*, 2001; Kolesnikova *et al.*, 2002; Noda *et al.*, 2002). Indeed, VP40 is able to produce virus-like particles (VLPs) when expressed in mammalian cells even in absence of other viral proteins (Harty *et al.*, 2000; Jasenosky *et al.*, 2001; Kolesnikova *et al.*, 2002; Noda *et al.*, 2002). Understanding the mechanism by which filoviruses assemble to form new virions, is tightly related to understanding the VP40 structure and function relationship with target lipids that may induce or stabilize VP40 oligomers.

VP40 forms a dimer (Bornholdt *et al.*, 2013; Oda *et al.*, 2016) with an amino-terminal domain (NTD) involved in dimerization and oligomerization and a carboxy-terminal domain (CTD) responsible for membrane binding (Bornholdt *et al.*, 2013; Oda *et al.*, 2016; Wijesinghe and Stahelin, 2016; Del Vecchio *et al.*, 2018), which may also function in oligomerization (Bornholdt *et al.*, 2013; Wijesinghe *et al.*, 2017). VP40 is a peripheral protein where mVP40 lipid binding was first speculated when the protein was shown to accumulate at intracellular membranes, mostly multivesicular bodies (MVB) and late endosomes early after its synthesis in cells (Kolesnikova *et al.*, 2002; Kolesnikova, Bamberg, *et al.*, 2004; Kolesnikova, Berghofer, *et al.*, 2004). Later, the critical role of anionic lipids, phosphatidylserine (PS) and phosphoinositides (PIP) for both mVP40 and eVP40 trafficking and interactions with the plasma membrane inner leaflet have been more well established (Ruigrok *et al.*, 2000; Adu-Gyamfi *et al.*, 2013, 2015; Johnson *et al.*, 2016; Oda *et al.*, 2016; Wijesinghe and Stahelin, 2016; Wijesinghe *et al.*, 2017).

Homo-oligomerization of the filovirus matrix protein is a key and required step for budding of virions (Nakai *et al.*, 2006; Adu-Gyamfi *et al.*, 2012b, 2015; Bornholdt *et al.*, 2013; Hilsch *et al.*, 2014; Freed, 2015; Johnson *et al.*, 2016). mVP40 and eVP40 are 34% identical in their amino acid sequence but only 16% identical in their CTDs, which gives rise to their different lipid binding selectivity. Differences in their CTDs may also contribute to differences in their oligomerization at the plasma membrane and within the cell. Indeed, mVP40 was previously described as forming large structures in cells (Timmins *et al.*, 2003; Liu *et al.*, 2010) and an octameric ring was observed when only the NTD (1-186aa) was purified (Timmins *et al.*, 2003). Timmins *et al* (Timmins *et al.*, 2003) hypothesized the paucity of distinct higher ordered mVP40 oligomeric structures was a result of the extremely high propensity of mVP40 (1-186) to oligomerize, indicated by the presence of extensive stacked rings (Timmins *et al.*, 2003). The same investigation successfully captured four distinct eVP40 oligomeric states, suggesting that mVP40 and eVP40 oligomerization may have fundamental differences (Timmins *et al.*, 2003). Furthermore, the dimeric and hexameric eVP40 crystal structures were resolved in 2013 lending significant insight to the origins of eVP40 lipid binding and oligomerization (Bornholdt *et al.*, 2013).

The current model of eVP40 oligomerization postulates that electrostatic interactions facilitate the disengagement of the eVP40 CTD from the NTD during matrix assembly. This disengagement sets up a conformational change which exposes two key residues within the NTD, Trp^95^ and Glu^160^, as part of an oligomeric interface. In 2016, the dimeric structure of mVP40 was resolved (Oda *et al.*, 2016) revealing a conserved Trp (Trp^83^) and Asn (Asn^148^) in mVP40 that align with eVP40-Trp^95^ and Glu^160^ (Fig. 1A, NTD panel), respectively. In a previous study, we reported that the Trp^83^ residue in particular was in a region that showed significant shielding during mVP40 membrane association using hydrogen-deuterium exchange-mass spectrometry (HDX-MS) analysis (Wijesinghe *et al.*, 2017), suggesting it may be important for mVP40 oligomerization following binding to anionic lipids. Furthermore, the previous work demonstrated a reduction of deuterium exchange at the CTD involving the residues Leu^226^ and Ser^229^ when mVP40 bound anionic membranes ((Wijesinghe *et al.*, 2017), Fig 1A). Notably, this region, dubbed α-helix 4 (α4 helix) just underlies the lipid binding surface and is distinct in residue composition and in structure when compared to eVP40. Therefore, we postulated two separate oligomerization interfaces within dimeric mVP40, one involving the CTD α4 helix as well as a conserved interface in the NTD as key regulators of mVP40 oligomerization (Wijesinghe *et al.*, 2017).

**Fig. 1.**
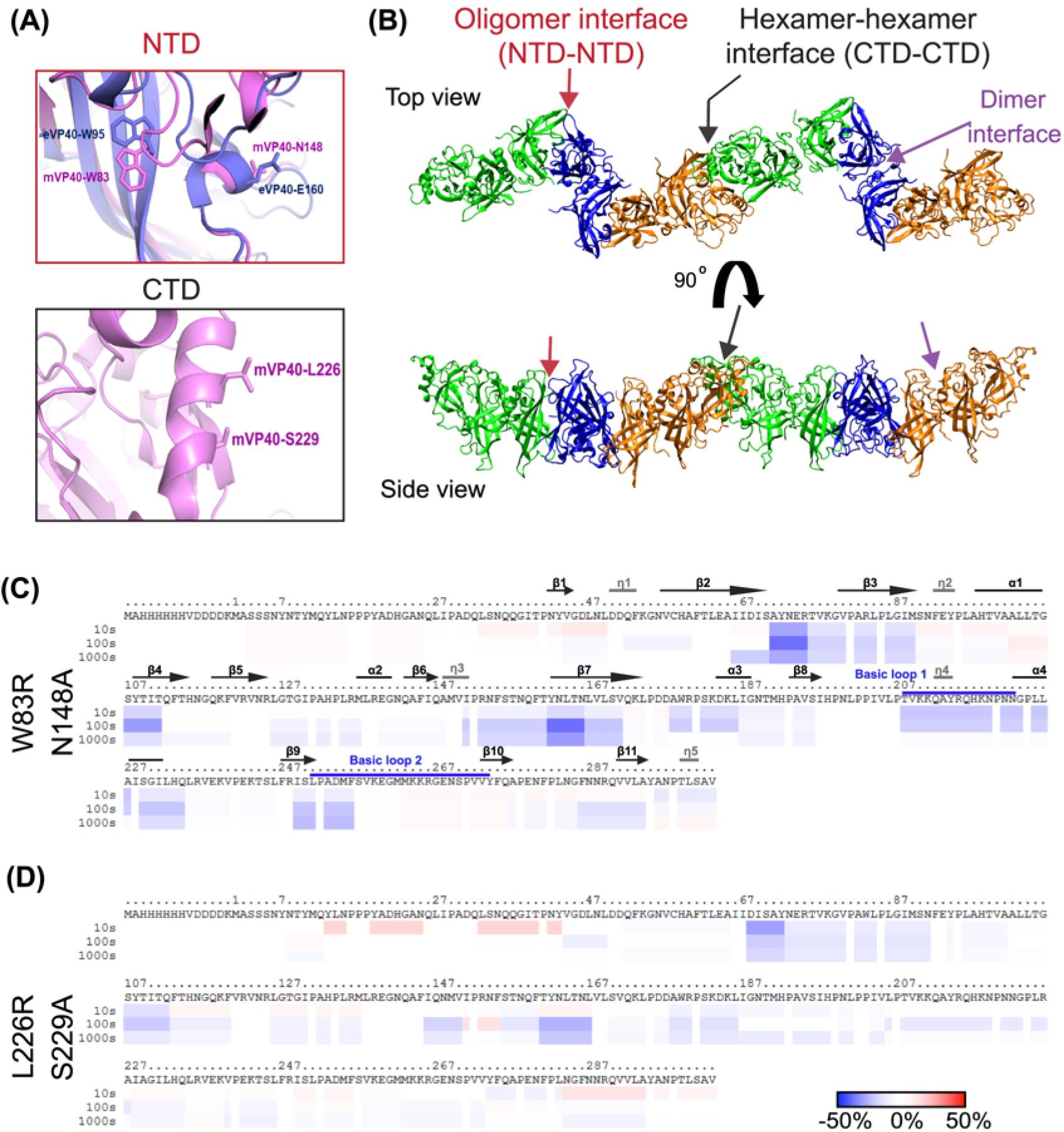
mVP40 potential oligomerization interfaces at NTD and CTD regions. **(A)** Zoomed in views of the structure of mVP40 at the NTD oligomer interface (upper panel) indicating Trp^83^ and Asn^148^ residues (pink) involved in the oligomerization with an overlay of Ebola virus VP40 (eVP40) structure with corresponding residues Trp^95^ and Glu^160^ (purple), and at the CTD interface (bottom panel) showing the potential residues Leu^226^ and Ser^229^ involved in hexamer-hexamer interactions. Modeled using PyMOL (mVP40 PDB ID: 5B0V) and (eVP40 PDB ID: 4LDB). **(B)** Top and side views of a mVP40 filament (two hexamers formed through the NTD-NTD interface, Fig. **S1**). **(C)** and **(D)** Ribbon maps of W83R/N148A and L226R/S229A mutants, respectively, indicating the difference in deuteration percentage of mVP40 in the presence of PS-containing liposomes. Each row corresponds to each time point collected (10 to 1000 s). Color coding: blue indicates the regions that exchange slower and red indicates the regions that exchange faster in the presence of liposomes.

To determine the mechanism of mVP40 oligomerization, we assessed different *in vitro* lipid binding assays with hydrogen/deuterium exchange mass spectroscopy (HDX-MS) analysis to study the effect of mutations at potential NTD and/or CTD oligomerization interfaces in mVP40 conformational changes upon binding membranes. Then, we conducted cellular studies to rationally investigate how the NTD and CTD oligomerization interfaces coordinate the matrix of mVP40 at the plasma membrane. Findings described here demonstrate that each oligomerization interface mutant displays a significant defect in VLP budding, consequence of impairment in overall and correct mVP40 trafficking and oligomerization at the plasma membrane.

## Results

### Effects of phospholipid membrane interaction on mVP40 oligomerization interface mutants

In order to better understand the origins of mVP40 oligomers in the absence of higher order mVP40 structural information, we constructed the mVP40 hexamer-hexamer interface using the eVP40 hexamer-hexamer interface as the template (PDB ID: 4LDD) and performed a 100-ns molecular dynamics simulation. Fig. 1A shows the section of the mVP40 filament composed of two hexamers next to each other involving CTD-CTD interactions (Fig. S1A). To test our hypothesis that both the conserved NTD and newly identified residues in the CTD are involved in mVP40 oligomerization, we first generated several mutant constructs. These included the NTD oligomerization interface mutant by W83R/N148A and a CTD oligomerization interface double mutant L226R/S229A. Size exclusion chromatography (SEC) of purified proteins indicated that all proteins formed dimers in solution (Fig. S2).

To dissect changes in mVP40 residue solvent accessibility and oligomerization in the absence and presence of membranes, HDX-MS experiments were performed on mVP40 mutants incubated with liposomes containing 45% phosphatidylserine (% molar ratio) as described previously (Wijesinghe *et al.*, 2017). In Fig. 1C, the ribbon map of the double mutant W83R/N148A indicates the differences in deuterium incorporation (%D) of the protein in presence of PS containing-liposomes compared to the protein alone. Overall, this double mutant showed little detectable changes in HD exchange pattern in both the NTD (from residue Met^1^ to Lys^47^) and CTD (from residue Met^263^ to Ala^284^). Similarly, residues Lys^96^-Gly^106^ on the helix α1 and residues Gln^112^-Phe^120^ on the β4-β5 strands exhibited slightly more rapid HD exchanges. Helix α1 is involved in the dimerization of mVP40 and it had been shown previously that the HD exchange at this region is slower in presence of anionic lipid-containing liposomes (Wijesinghe *et al.*, 2017). The HDX-MS profile of W83R/N148A also showed an increase of HD exchange at the β6 strand (residues Glu^140^-Gln^146^) as well as in the region Met^261^ to Gln^276^, which is in basic loop 2 and the β10 strand. Oda et al. (Oda *et al.*, 2016) showed that residues in this region are involved in the efficient binding of mVP40 to PS-containing liposomes. All together, these results suggest that the residues Trp^83^ and Asn^148^ are involved in the formation of oligomers which shields these specific regions from exposure to aqueous environment resulting in slow deuterium incorporation/exchange rates upon binding to PS-containing lipid vesicles. Furthermore, the double mutant W83R/N148A exhibited an intermediate change in deuteration level compared to wild type mVP40 (WT-mVP40) in presence or absence of zwitterionic phospholipid (Fig. S1B adapted from (Wijesinghe *et al.*, 2017)).

Next, we analyzed the solvent accessibility of the CTD double mutant L226R/S229A upon binding to PS-containing lipid vesicles. Similar to W83R/N148A, L226R/S229A exhibited an overall increase of the HD exchange profile compared to WT-mVP40 (Fig. S1B), except in the region including residues Ile^88^-Asn^91^. Further, no changes of the deuteration level of the β6 strand (residues Glu^140^-Phe^145^) were observed for L226R/S229R compared to WT-mVP40, which showed very slow HD exchange in presence of PS-containing vesicles within the same region (Wijesinghe *et al.*, 2017). As mentioned above, L226R/S229A-mVP40 showed a faster HD exchange profile than WT-mVP40, including the following regions: i) in the NTD from residue Tyr^13^ to Tyr^44^ (which contains the β1 strand), residues Glu^73^-Gly^87^ (unstructured loop between β2-β3 strands and the N-terminus of β3 strand), Phe^113^-Phe^120^ (β4-β5 unstructured loop) and residues Ile^146^-Asp^177^ (which includes the unstructured loop between helix η3-β7 strand and the entire β7 strand (Fig. 1D); ii) in the CTD of mVP40, the entire region including residues Thr^208^-Lys^264^, which contains helices η4 and α4, the unstructured loops between these two helices, β9 strand as well as the unstructured loop β9-β10 harboring basic loop 2, and finally the region containing the β11 strand until the C-terminus (Fig. 1D). Altogether, this analysis suggests that mutation of the hypothesized CTD oligomerization interface reduces oligomerization of mVP40 in presence of PS-containing vesicles resulting in an exposure of residues at or close to the CTD of the protein.

### Mutations of key residues in mVP40-NTD or CTD oligomerization interfaces alter mVP40 plasma membrane localization

To investigate the role of the NTD and CTD oligomerization interfaces of mVP40 on the protein trafficking and binding to the plasma membrane, we performed live cell imaging of EGFP tagged WT mVP40, single mutants of each oligomerization interface, W83R and L226R, as well as the double mutants W83R/N148A and L226R/S229A (Fig. 2A-B). In agreement with previous investigation, WT-mVP40 primarily associates with the plasma membrane (Fig. 2A). Confocal imaging illustrated the ability of the mutant EGFP-W83R-mVP40 to traffic and localize to the plasma membrane (Fig. 2A), to a level comparable to WT-mVP40 (Fig. 2B). Additionally, W83R did exhibit elongated structures at the plasma membrane similar to WT, which corresponds to assembled VLPs. The double mutant EGFP-W83R/N148A-mVP40 exhibited similar membrane localization deficiency to monomeric mutant T105R (Fig. 2A-B). This result is consistent with previous data described in Oda et al (2016) and Koehler et al (2018). However, unlike WT-mVP40, no significant intracellular aggregations were observed in W83R/N148A expressing cells 14 hours post-transfection (Fig. 2A). Co-expression of mVP40 and mutations with mGP increased the plasma membrane localization of W83R/N148A by 11% (Fig. 2A-B). However, despite this modest increase in plasma membrane localization, no elongated tubulations were detected on the surface of transfected cells (Fig. 2A), which are abundant on cells expressing WT-mVP40 (in absence or presence of mGP, see Fig. 2A top left panel). These observations may indicate the requirement of an interaction with both Trp^83^ and Asn^148^ residues within the NTD oligomerization interface for efficient membrane localization of the protein and assembly into VLPs.

**Fig. 2.**
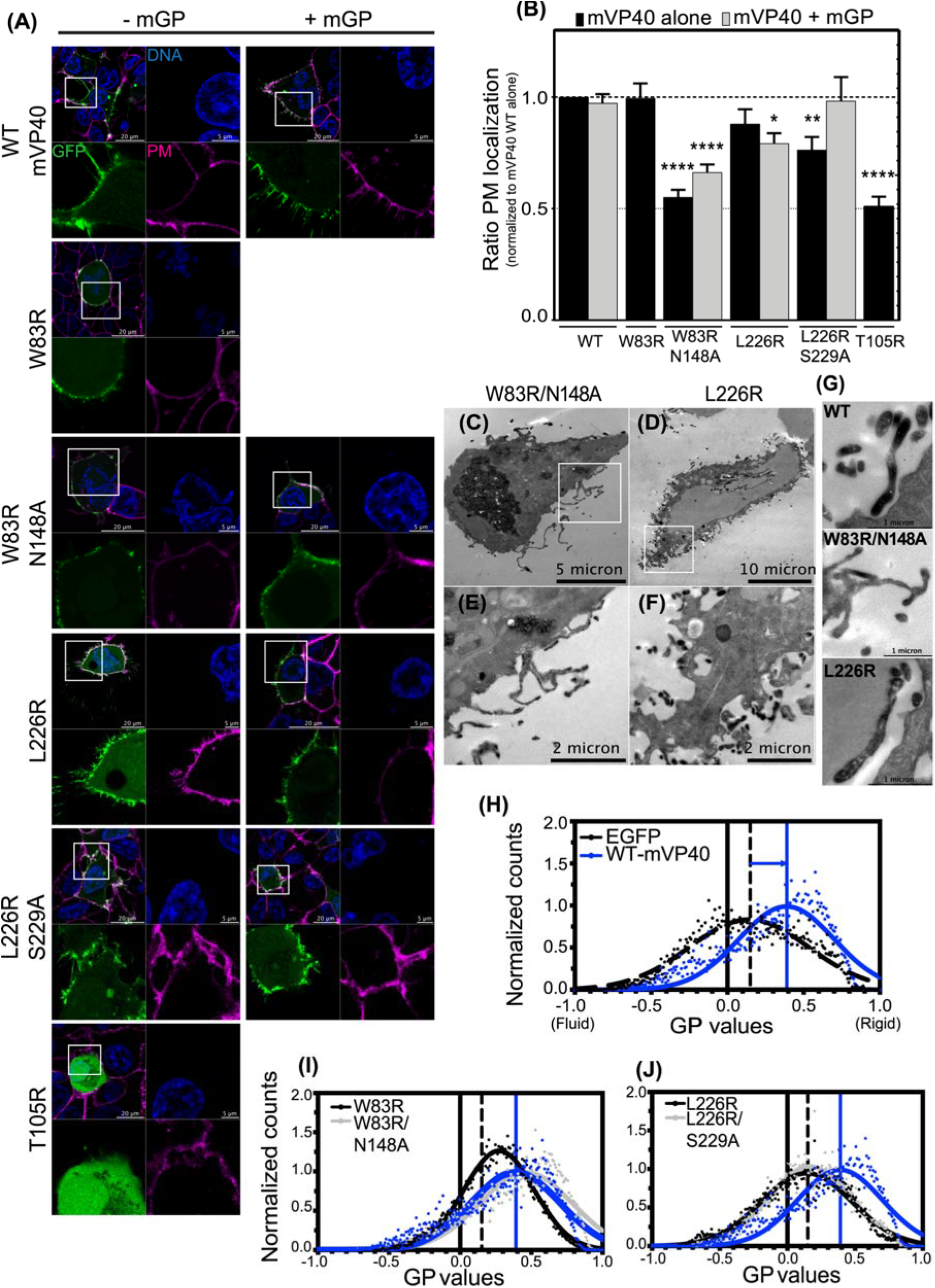
NTD and CTD oligomerization interfaces required for efficient mVP40 trafficking to the plasma membrane. **(A)** Confocal live images of cells expressing EGFP-constructs (green) +/− glycoprotein mGP, stained for DNA (blue) and plasma membrane (PM, pink). **(B)** Ratio of PM retention **from A** quantified by calculating the integrated density of pixels at PM to total pixels within the cell and normalized to WT. Data are represented as averages ± S.E.M of three independent means. Statistical analysis was performed using one-way ANOVA with multiple comparison Holm-Sidak tests, (* *p*=0.01, ** *p*=0.001, **** *p*< 0.0001). **(C)** and **(D)** are representative TEM micrographs of HEK293 cells co-expressing GBP-APEX2 and EGFP-mVP40 W83R/N148A and L226R, respectively. **(E)** and **(F)** Zoomed insets in **(C)** and **(D)** respectively. **(G)** TEM micrographs of potential VLPs at cell surfaces when expressing EGFP-mVP40 indicated constructs. Experimental and fitted normalized general polarization (GP) distribution curves of laurdan dye across PM of HEK293 cells with EGFP (black dashed line), **(H)**mutants of NTD **(I)** and CTD **(J)** oligomerization interfaces, compared to WT (blue line). GP values range from − 1 (very fluid lipid domains) to + 1 (very rigid lipid domains). The fitting procedure was performed using a non-linear Gaussian curve.

In contrast, the single mutant EGFP-L226R-mVP40 showed a non-significant decrease in plasma membrane localization in both the presence and absence of mGP co-expression (Fig. 2A-B). However, the double mutant EGFP-L226R/S229A-mVP40 had a more pronounced and significant reduction in localization at the plasma membrane compared to WT-mVP40, (25% reduction) (Fig. 2A-B). These observations may suggest collaborative interactions at the CTD between L226 and S229 to ensure normal plasma membrane enrichment of mVP40. Both the L226R and L226R/S229A mutants were still able to form filamentous protrusions at the plasma membrane. Furthermore, co-expressing the CTD oligomerization interface double mutant L226R/S229A with mGP appeared to fully recover the wild type phenotype (Fig. 2A-B). Taken together, these results indicate that the residues involved in NTD oligomerization interface are essential to matrix assembly at the plasma membrane for the elongation of VLPs while the CTD oligomerization interfaces may be required for efficient trafficking and binding of mVP40 to the plasma membrane of mammalian cells. This was further supported by the lack of reduction in deuterium exchange for L226R/S229A in regions of membrane binding previously mapped for mVP40 (Oda *et al.*, 2016; Wijesinghe *et al.*, 2017). As a control, we also analyzed the monomeric mVP40 mutant T105R that had been shown to exhibit diffused signal in the cytosol (Oda *et al.*, 2016). As expected, EGFP-T105R-mVP40 failed to translocate to the plasma membrane (Fig. 2A-B).

### mVP40-NTD oligomerization interface mutations abolish the ultrastructure of VLP at the plasma membrane

For a clearer understanding of the role of each oligomerization interface in mVP40 multimerization and assembly at the host plasma membrane, we performed transmission electron microscopy (TEM) imaging of the W83R/N148A double mutant and L226R single mutant. We chose these two mutants because of their altered phenotypes in cells and because we observed highly similar VLP structures in the L226R and L226R/S229A expressing cells from live confocal imaging (Fig 2A, left panel). To ensure we only evaluated cells expressing mVP40, we performed a novel ascorbate peroxidase-tagging (APEX) TEM method which utilizes the co-expression of EGFP-tagged proteins with GFP binding protein (GBP) fused to APEX2 (GBP-APEX2) (Ariotti *et al.*, 2015). Upon co-expression of GBP-APEX2 with GFP-mVP40 proteins, GBP-APEX binds to the EGFP tag on mVP40. During TEM processing, APEX2 catalyzes the conversion of diaminobenzidine (DAB) into a precipitate that deposits at the site of the GBP-APEX2:GFP interaction. Following chemical fixation, the precipitate allows a specific and localized signal of EGFP-mVP40 localization with high contrast for TEM imaging.

First, we tested the ability of WT-mVP40 to translocate to the plasma membrane and to form the typical elongated structure of VLPs in cells co-transfected with EGFP-WT-mVP40 and GBP-APEX2. As shown in Fig. S3B and Fig. S3C, the co-expression of the two constructs resulted in normal VLP protrusions from the plasma membrane. To validate this observation, we compared post-stained cells expressing EGFP-WT-mVP40 alone (Fig. S3A, S3D) to cells expressing both EGFP-WT-mVP40 and GBP-APEX2 (Fig S3B, S3E) and found that VLP structures between both conditions were morphologically indistinguishable. We also assessed if post-staining APEX2 expressing cells could enrich the contrast detected for TEM, as post-stain enhances membrane staining for organelle identification, therefore we compared cells expressing GBP-APEX2 and EGFP-WT-mVP40 with and without post-stain. Cells that did not receive post-stain yielded superior APEX2 signal at the membrane of the cell and the VLP membranes where mVP40 is enriched (Fig. S3B, S3E) compared to post-stained APEX cells (Fig. S3C, S3F). The post-stain appeared to introduce artifacts at the plasma membrane which could alter our observations and analysis. Therefore, we decided to continue our investigations of mVP40 mutants co-expressed with GBP-APEX2 without any post-stain processing.

Fig. 2C and 2D are representative micrographs of cells co-expressing GBP-APEX2 and EGFP-W83R/N148A or EGFP-L226R-mVP40, respectively. The APEX2 signal from EGFP-W83R/N148A mutant was more distributed within the cell (Fig. 2C) with some distinct puncta at the membrane and across some tubulations (Fig. 2E) that may correspond to accumulated mVP40. However, the mutant did not show characteristic VLP structures found at the plasma membrane of WT-mVP40 (Fig. 2G top image) but instead moderate APEX2 signal in ruffled membranes (Fig. 2G middle image). On the other hand, APEX2 signal from EGFP-L226R mutant was located at the cell periphery (Fig. 2D) and was detected inside VLP structures (Fig. 2F). Also, no major defect in the ultrastructure of the VLPs was observed in this mutant although their abundancy at the plasma membrane was possibly different (Fig. 2G bottom image). Taken together, this analysis corroborated our confocal imaging results where the mutations of residues within the NTD oligomerization interface impaired the accumulation of mVP40 at the plasma membrane and the proteins ability to assemble and form VLPs unlike the mutation within the CTD oligomerization interface.

### Plasma membrane fluidity exhibits a different profile upon binding of mVP40 variants

We hypothesized that membrane fluidity changes may be important for proper mVP40 matrix assembly and virus particle elongation and that mVP40 oligomerization may facilitate this process. To test this, we employed a laurdan fluidity imaging assay of cells expressing the different EGFP-mVP40 variants (or EGFP plasmid as a control) (Fig. S4). Laurdan is a fluorescent hydrophobic probe that penetrates cell membranes and aligns parallel to the phospholipid tails (Bagatolli *et al.*, 2003). In ordered or rigid membranes with a highly hydrophobic environment, the probe has a peak emission wavelength of ◻ 440 nm. However, in fluid membranes water molecules adjacent to the glycerol backbone induce dipolar relaxation of laurdan, resulting in a spectral shift in the emission wavelength to ◻ 500 nm (Gaus *et al.*, 2006). Changes in membrane fluidity can then be measured by a normalized ratio of the two emission regions, and is called the general polarization (GP) index (which ranges between −1 and 1, for fluid to rigid membranes, respectively) (Bagatolli *et al.*, 2003).

The analysis of laurdan fluorescence was performed under a two-photon confocal microscope and we focused the analysis on cells with the largest enrichment of mVP40 at the plasma membrane (Fig. S4, bottom panels). The GP index shifted from 0.15 at the plasma membrane of HEK293 cells expressing EGFP to ◻ 0.4 at the plasma membrane of EGFP-WT-mVP40 expressing cells (Fig. 2H). This observation suggests that the binding and assembly of mVP40 at the plasma membrane increases membrane rigidity. Next, to investigate the role of different oligomerization processes during the matrix assembly at the plasma membrane on its fluidity, we analyzed the GP index of NTD and CTD oligomerization interface mutants compared to WT-mVP40 (Fig. 2I, 2J). Expression of W83R in cells did increase membrane rigidity compared to EGFP alone (GP index ~0.25 in W83R expressing cells), albeit to a lesser extent than WT (Fig. 2I). Interestingly, the double mutant W83R/N148A had a comparable effect as WT on the plasma membrane rigidity, with a GP index of ◻ 0.45 in W83R/N148A expressing cells (Fig. 2I). Conversely, L226R and L226R/S229A mutants exhibited exactly the same GP (◻ 0.15) as EGFP expressing cells (Fig. 2J), indicating their association with membranes does not change membrane fluidity. All together, these data suggest that the membrane rigidity observed in the wild type is a result of CTD-CTD oligomerization of the virus matrix at the plasma membrane.

### Mutation of mVP40 at hypothesized oligomerization NTD interfaces drastically reduced mVP40 oligomerization in cells

To confirm a defect in the matrix assembly upon NTD and CTD oligomerization interface mutations, we assessed cellular mVP40 oligomerization analysis through Number & Brightness (N&B) analysis. N&B is a method used to analyze the assembly state of proteins in real-time based on the variance of the intensity within single pixels over time (Digman *et al.*, 2008). Moreover, this technique has been used to evaluate viral matrix protein oligomerization (Adu-Gyamfi *et al.*, 2015; Johnson *et al.*, 2016).

To determine the brightness value for a monomer, GFP was expressed in HEK293 cells. To determine the brightness value of higher ordered oligomeric states of GFP-mVP40 constructs expressed in HEK293 cells, multiples of the EGFP monomer brightness value were extrapolated to the corresponding oligomeric states. Pixel intensities correlating to monomer-hexamer (red), hexamer-12mer (green), 12mer-24mer (blue), and >24mer (pink) oligomeric states of mVP40 were analyzed, mapped onto the original composite image of the cell and plotted as a percent of total pixels in the image (See Fig. S5A). The oligomerization profile of EGFP-WT-mVP40 revealed the largest population of mVP40 was in the monomer-hexamer assembly state (~52.62% total pixels, Fig. 3A, Table S2, Fig. S5A). Importantly, each higher ordered oligomeric state was roughly equally represented (from ~13% to 19.1% total pixels, Fig. 3A, Table S2, Fig. S5A).

**Fig. 3.**
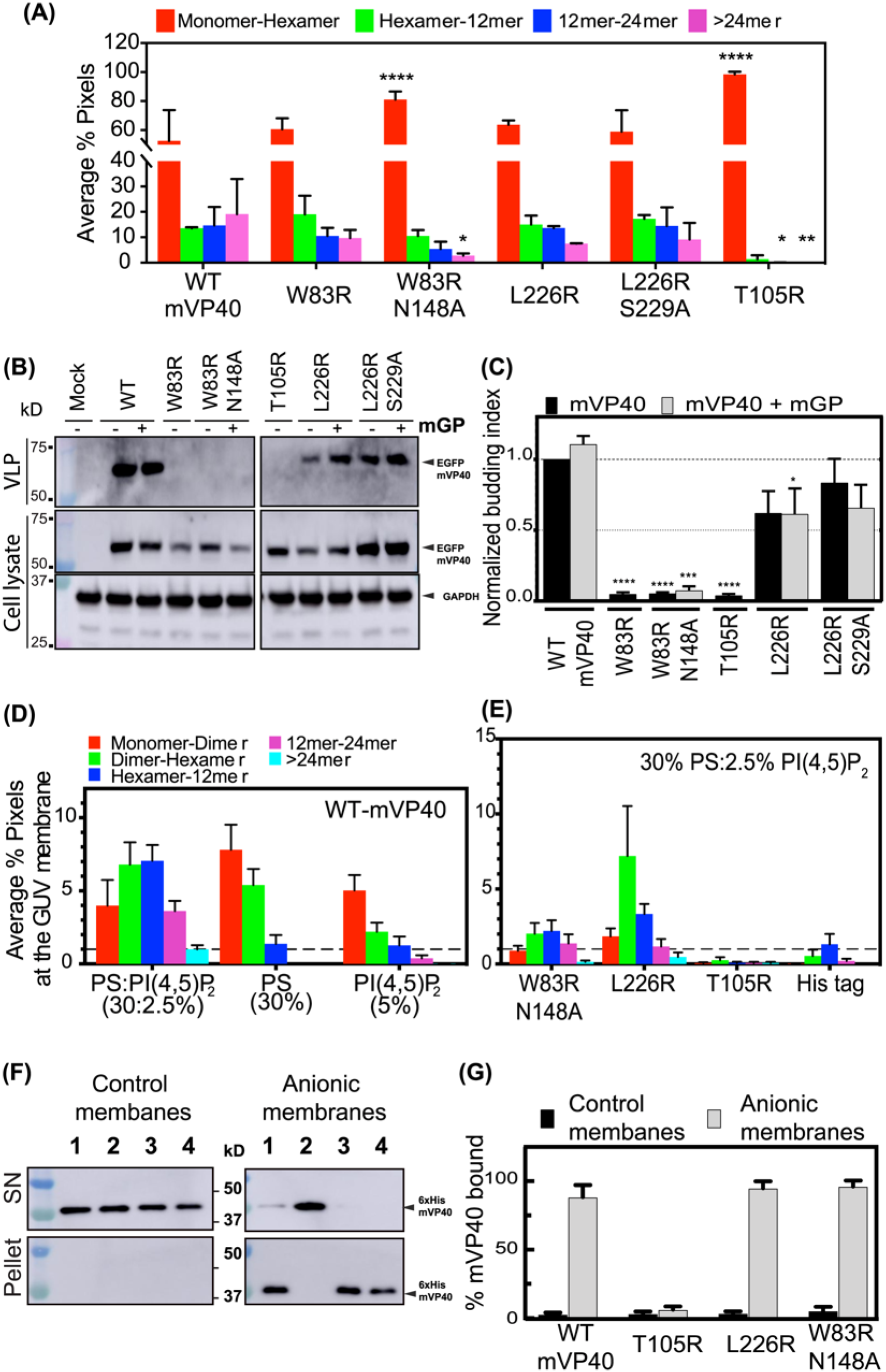
Cellular and *in vitro* oligomerization are impaired in NTD and CTD interface mutants reducing VLP budding. **(A)** Average % pixels with each estimated oligomerization form from N&B analysis performed 24 h.p.t of HEK293 cells with EGFP-mVP40 constructs. Functional budding assays were performed to assess the capacity of WT-mVP40 and mutants to produce VLPs. **(B)** Representative Western blot assays performed on VLPs (top panel) and cell samples (middle and bottom panels) from cells 24 h.p.t in the presence and absence of glycoprotein. **(C)** Quantification of the budding index for each mVP40 protein (normalized to mVP40 WT) was determined by densitometry analysis. **(D)** Plotted average % pixel from N&B analysis of WT-mVP40 enriched at GUV membranes indicating the oligomerization profile of mVP40. **(E)** Oligomerization profiles of W83R/N148A, L226R, the monomeric mutant T105R and His-tag alone at the PS:PI(4,5)P_2_-containing membranes. **(F)** binding efficiency of WT-mVP40 (lane 1) and mutants (lane 2: T105R, lane 3: L226R, lane 4: W83R/N148A) to anionic membrane (30% PS:2.5% PI(4,5)P_2_) assessed by liposome sedimentation assay and quantified in **(G)**. Values are reported as mean ± S.D **(A, G)** or ± S.E.M **(D, E)** of three independent means. One-way ANOVA **(C)** or two-way ANOVA **(G)** with multiple comparisons were performed. (\**p*<0.05, \*\*\**p*<0.0005, \*\*\*\**p*<0.0001).

Analysis of the oligomerization profile of each EGFP-mVP40 mutant differed from the WT oligomerization profile. In the NTD oligomerization interface mutant W83R, we noted a reduction of ~10% in large oligomers >24mer (from 19.1% to 9.68%) with a non-significant but proportional increase of 8% in monomer-hexamer (from 52.62% to 60.7%) and ~ 6% in hexamer-12mer (from 13.58% to 19.04%) (Fig. 3A, Table S2, Fig. S5A). A similar but more significant pattern was observed for the double mutant W83R/N148A, where a significant increase in monomer-hexamer was observed (~29% increase from 52.62% to 81.15%) concomitantly with a notable decrease of ~16% in oligomers >24mer (decreased from 19.1% to 2.8%) (Fig. 3A, Table S2, Fig. S5A). Given these findings, this analysis demonstrated that the mutants have an impaired ability to form large oligomers and accumulated at the plasma membrane in small oligomers. Unfortunately, this experiment could not provide further information on the ability of the mutants to form a hexamer through NTD-NTD oligomerization. In contrast, CTD oligomerization interface mutants did not exhibit a drastic change in their oligomerization profile compared to WT except for a slight decrease in oligomers >24mer (~12% and ~10% reduction for L226R and L226R/S229A, respectively) and modest increase of monomer-hexameric structures (~11% and ~ 7.5% increase for L226R and L226R/S229A, respectively) (Fig. 3A, Table S2, Fig. S5A). In brief, CTD oligomerization interface mutants have a smaller effect on the ability of the protein to form large oligomers >24mer that may involve other residues and may have a compensatory effect. The monomeric and nonfunctional T105R mutant was used as control and did not show any detectable oligomerization (Fig. 3A, Table S2, Fig. S5A). Altogether, these results support our hypothesis of a potential oligomerization interface in the NTD required for efficient mVP40 matrix assembly and virus budding.

### NTD oligomerization deficient mVP40 mutants fail to produce VLP

To understand the functional significance of mVP40 oligomerization deficient mutants, functional budding assays of HEK293 cells expressing EGFP-mVP40 variants were performed. We hypothesized that mVP40 mutants with aberrant oligomerization would fail to produce VLPs. Additionally, an interaction between mGP and mVP40 has been previously reported (Kolesnikova *et al.*, 2007), therefore, co-expression of mGP and mVP40 was performed for some of the functional budding assays. Robust VLP production was observed for cells expressing WT-mVP40, with a slight increase in VLP production when WT-mVP40 was co-expressed with mGP (Fig. 3B, 3C). Both NTD oligomerization interface mutants lost their ability to release VLPs as described previously in (Oda *et al.*, 2016) and (Koehler *et al.*, 2018), even in the presence of mGP (Fig. 3B, 3C). These results demonstrated that despite the ability of W83R and W83R/N148A mutants to bind and form small oligomers at the plasma membrane, their deficient ability to form large oligomers results in an inability to release VLPs. On the other hand, CTD oligomerization interface mutants had a reduction of VLP release of ~40 % for L226R (in the presence and absence of mGP), and of ~25% for the double mutant L226R/S229A in the absence of mGP. Interestingly, co-expression of L226R/S229A with mGP resulted in an even more profound reduction in VLP production compared to WT (~40% reduction) than when L226R/S229A was expressed alone (Fig. 3B, 3C). The APEX2 TEM analysis of VLP structures at the plasma membrane of the L226R mutant did not show a significant morphological defect (Fig. 2F, 2G bottom image), however, the functional budding assay suggests a defect in efficient scission from the plasma membrane to form VLPs. Taken together, these observations highlight the importance of the L226 and S229 residues in the CTD oligomerization interface to ensure a functional mVP40, despite the ability of these mutants to multimerize and form a matrix and elongate at the plasma membrane, albeit to a lesser extent than WT. This also underscores the importance of CTD-CTD mediated changes in membrane rigidity, which may be an important step in the proper matrix layer formation for effective scission. The monomeric nonfunctional T105R mutant was used as control of budding deficiency and failed to produce VLPs (Fig. 3B, 3C).

### Oligomerization profiles of wild type mVP40 at the surface of GUVs depends on the anionic lipid compositions

The oligomerization profile in cells of mVP40 variants did not provide adequate details concerning the profile of small oligomers at the plasma membrane. This may be due to non-bound proteins in the cytosol that may make the distinction of pixel intensities correlating to monomer-dimer from the one correlating to dimer-hexamer, in addition to intracellular factors that could also promote protein oligomerization. In order to address this point, we performed N&B analysis on purified 6xHis-mVP40 proteins incubated with giant unilamellar vesicles (GUVs) and this analysis required a fluorescent protein. For this purpose, we first attempted to conjugate dimeric 6xHis-mVP40 through its unique cysteine residue (Cys^58^) using maleimide-AlexaFluor conjugated dye. However, we were unable to conjugate efficiently mVP40 despite multiple attempts (data not shown), which is likely due to the low structural accessibility of this residue to solvent. Alternatively, we used a (Ni)-NTA-Atto550 conjugate probe that is specific for poly-histidine tags with minimal cross reactivity (You and Piehler, 2014).

Previously, Wijesinghe & Stahelin (Wijesinghe and Stahelin, 2016) demonstrated that mVP40 associated non-specifically with the anionic lipids within the plasma membrane (e.g. PS and PIPs). Here, we aimed to understand the oligomerization profile of mVP40 during virus matrix assembly at the plasma membrane of infected cells, the building block of the virus particles. For this reason, we used the well-established giant unilamellar vesicle (GUV) assay, which allows tailored lipid compositions with the ability to incorporate small amounts of fluorescent lipids for visualization. Thus, using the GUVs, we are able to selectively determine binding and oligomerization of mVP40 in the presence of PS, PI(4,5)P_2_ and/or both. Confocal imaging was performed to ensure the Ni-NTA-Atto550/His-WT-mVP40 bound efficiently to the GUVs (Fig S5B, composite panels). Then, we determined the brightness value for the monomeric Ni-NTA-Atto550 dye. Thus, we performed confocal microscopy imaging of WT-mVP40 proteins incubated with different GUV compositions followed by N&B analysis (Fig. 3D-3E; brightness plots in Fig. S5B). Only pixels detected at mVP40-enriched GUV membranes were analyzed and normalized to the total amount of pixels detected to estimate the oligomeric distribution across the *in vitro* membrane.

This analysis demonstrated for the first time that mVP40 protein oligomerization profiles depend on the lipid composition of the membrane. Indeed, WT-mVP40 is able to bind PS:PI(4,5)P_2_ containing GUVs, where ~25% of the total pixel counts corresponded to membrane bound mVP40 (Fig 3D) and more than 60% of the bound protein formed approximately equal population of dimer-hexamer and hexamer-12mer at the vesicle membrane (30.29 and 31.44%, respectively, Table S3). For the remaining fraction of mVP40 membrane bound, ~17.7% was monomer-dimer and ~16% was of 12mer-24mer. Finally, 4.43 % of total bound mVP40 were very large oligomers, >24mer. Furthermore, in PS-containing GUVs, WT-mVP40 was detected mostly as small oligomers with an abundancy of monomer-dimer (~53% total bound protein, Table S3) and dimer-hexamer (~37% total bound protein). However, only a small population of hexamer-12mer was detected (~9.4%) and no larger oligomers could be detected (>12mer) without PI(4,5)P_2_ in the GUVs. This first analysis suggests that both PS and PI(4,5)P_2_ are required for mVP40 to form larger oligomers and assemble the viral matrix. Moreover, in PI(4,5)P_2_ containing GUVs, a small population of pixels at the membrane of the GUV were detected, this may explain the low abundance of negative charge at the surface of the membrane (20% of total charge) compared to the previous liposomes compositions, 50% and 30% respectively (Wijesinghe and Stahelin, 2016). Overall, ◻ 56.5% of the bound protein was monomer-dimer, 24.72% as dimer-hexamer, 14.25% as hexamer-12mer and 4.25% 12mer-24mer while no >24mer could be detected (Fig. 6A, Table S3). This result indicates that PI(4,5)P_2_ may promote mVP40 to form larger oligomers (over than 12mer), which requires the presence of PS in the GUVs. To summarize, this analysis demonstrated a different oligomerization profile of mVP40 depending on the lipid composition of vesicle membrane, where both PS and PI(4,5)P_2_ are required for large VP40 oligomers suggesting that PS is sufficient to promote small VP40 oligomers such as hexamers while PI(4,5)P_2_ is likely involved in promoting or stabilizing hexamer-hexamer interactions.

### In vitro oligomerization of mVP40 is altered upon mutation of key residues

To investigate the effect of NTD and CTD oligomerization interface mutations on *in vitro* oligomerization, we performed N&B analysis using 6xHis-W83R/N148-mVP40 and 6xHis-L226R-mVP40 purified proteins. We confirmed by size exclusion that these two mutants formed the dimer (Fig. S2) indicating that the mutations had no effect on the dimerization of the protein. Also, we decided to continue our investigations using only these mutants as W83R/N148A had a more profound phenotype in cells compared to W83R and due to L226R and L226R/S229R displaying similar phenotypes in cells (aside from impaired plasma membrane binding of L226R/S229R). First, we compared the oligomerization profiles of both W83R/N148A and L226R on GUVs that contained both PS and PI(4,5)P_2_ (Fig. 3E, Table S3). W83R/N148A showed efficient binding to the GUV membranes; however large oligomer formation was significantly reduced (Fig. 3E, Table S3). In this analysis, a small population of pixels was detected at mVP40-enriched GUV membranes (6.62 % total pixels). Because we focused this analysis on mVP40 enriched GUVs, it is important to note that this small population of pixels detected did not suggest a defect of binding to the GUV membrane. The distribution of mVP40 consisted of 13.31% of the total bound protein profile as monomer-dimer, 30.29 % dimer-hexamer, 33.31% hexamer-12mer, 20.7% for 12-24mers, and 2.17% for oligomers >24mer, of total bound protein (Fig. 3E, Table S3). The major differences between this NTD mutant and the WT oligomerization profile is the increase of hexamer-12mer and 12mer-24mer population in the W83R/N148A mutant compared to the WT (Fig 3D, 3E, Table S3). Additionally, L226R displayed a unique oligomerization profile where the most abundant structures were dimer-hexamer, with over 51% of total bound protein (14.02% total pixels) at the membrane of GUV containing PS:PI(4,5)P_2_ (Fig. 3E, Table S3). The other oligomers, in contrast to WT and the NTD mutant, exhibited a decrease in their abundancy with 13.15% monomer-dimer, 23.77% hexamer-12mer and 8.37% of 12mer -24mer. This result strongly supports our hypothesis that the α4 helix and residue Leu^226^ plays a critical role in oligomerization by facilitating CTD-CTD interaction to expand the matrix from a hexamer to larger filaments *in vitro*.

We next extended our investigations into the role of specific lipids in facilitating mVP40 oligomerization at both the NTD and CTD oligomerization interfaces by performing N&B using W83R/N148A and L226R on GUVs that contained only PS or only PI(4,5)P_2_ (Fig. S5C, S5D). First, we observed that both mutants displayed a high abundance at the PS-containing membrane as a monomer-dimer, 79.49% and 69.07% of total bound protein, respectively. In contrast, a reduction of other oligomers was observed for W83R/N148A and L226R with dimer-hexamer (18.78% and 25.11%, respectively) and hexamer-12mer (1.73% and 5.72%, respectively) (Fig. S5C, Table S3). Neither WT-mVP40 or either mutant were able to form larger oligomers on PS only GUVs (Fig. 3D, Table S3, Fig. S5C); however, the oligomerization profile of W83R/N148A was notably defective compared to both WT and L226R on PS only GUVs.

We next performed N&B using GUVs with only PI(4,5)P_2_. Using PI(4,5)P_2_ vesicles we demonstrated that the W83R/N148A mutant had a comparable oligomerization profile to WT, with a small increase of hexamer-12mer population from 14.25% in the WT to 16.25% in the mutant. W83R/N148A also exhibited a depletion of the >24mer population with less than 1% of total bound protein (Fig. S5D, Table S3). The single mutant, L226R, showed a high enrichment at the membrane of PI(4,5)P_2_-containing GUVs, where ◻ 22% of total pixels corresponded to bound mVP40, compared to 8.89% for the WT and 11.73% for W83R/N148A, and a higher population of dimer-hexamer with 28.87% of total bound protein (24.72% and 26.36% for WT and W83R/N148A, respectively) (Fig. S5D, Table S3). The other oligomers detected for the L226R mutant were 45.49% of monomer-dimer, 17.45% of hexamer-12mer, 6.04% of 12mer-24mer and 2.12% of >24mer of total bound protein (Fig. S5D, Table S3). The monomeric mVP40 mutant T105R and 6xHis-tag were used as controls for no binding and oligomerization on GUV membranes (Fig. S5C, S5D, Table S3). Taken together, we demonstrated that both W83R/N148A and L226R mutants exhibited oligomerization profiles that are consistent with the role of the mutated residues in mVP40 matrix assembly, where CTD oligomerization interface mutant L226R displayed an accumulation of dimer-hexamer population in both PI(4,5)P_2_ and PS:PI(4,5)P_2_-containing vesicles; on the other hand, NTD oligomerization interface double mutant W83R/N148A accumulated mostly as monomer-dimer at PS-containing membranes with a deficiency to form other oligomers.

### Association with anionic lipids is not altered in mVP40 oligomerization interface mutants

To assess the effect of NTD and CTD interface mutations on the ability of mVP40 to bind PS:PI(4,5)P_2_-containing membranes *in vitro*, a liposome sedimentation assay was performed. A representative Western blot is shown in Fig. 3F and quantified results from densitometry analysis are shown in Fig. 3G. LUVs were prepared containing either no anionic lipids (control membranes) or with 30% PS and 2.5% PI(4,5)P_2_ (anionic membranes). This assay showed a clear ability of all proteins to efficiently bind anionic membranes with no detectable binding to control membranes (Fig. 3F, 3G). The monomeric T105R mutant was used as a control and lacked detectable binding to membranes (Fig. 3F, 3G) demonstrating the necessity of the intact dimer in binding anionic membranes as previously reported (Oda *et al.*, 2016). This suggests that both NTD and CTD oligomerization interfaces are not involved in mVP40 binding to anionic phospholipid-containing membranes and that observations from orthogonal experiments are not a result of an inability of the protein to associate with PS or PI(4,5)P_2_ containing membranes or the plasma membrane.

### NTD/CTD oligomerization interfaces triple mutant displayed a unique profile

To deepen our understanding of the oligomerization process of mVP40 and the role of both NTD and CTD oligomerization interfaces in this process as well as in the viral matrix assembly, we generated a 6xHis-mVP40 triple mutant of both the NTD and CTD oligomerization interfaces ((W83R, N148A and L226R (WNL-mVP40)). SEC of purified protein indicated that the triple mutant formed a dimer in solution (Fig. S2). HDX-MS analysis was performed with membranes as described above (Fig. 4A) and demonstrated that WNL displayed an overall decrease of HD exchange compared to WT-mVP40 except for four regions: C-terminal region of β2 strand (Ile^66^-Ser^70^), β6 strand (residues Glu^140^-Phe^145^), N-terminal region of β7 strand (residues Leu^167^-Val^171^) and basic loop-2 with the β10 strand (residues Lys^265^-Gln^276^). However, the two last regions had an increased rate of HD exchange at longer time points. On the other hand, some regions showed a slower HD exchange than WT, which included residues Ala^71^-Arg^75^ in the loop region between the β2 and β3 strands, residues Tyr^162^-Asn^166^ within the β7 strand, the residues constituting helix α3 (Lys^183^-Ile^187^), residues Ile^249^-Val^259^ found in the β9 strand and the N-terminal region of the basic loop-2 and residues Asn^280^-Tyr^295^ in the unstructured loop between the β10 and β11 strand (Fig. 4A). Other regions displaying low HD exchange at longer time points included residues in unstructured loops, Tyr^157^-Asn^166^ (unstructured loop between η3 and N-terminus of β7 strand), Tyr^208^-Arg^226^ (unstructured loop between η4 and helix α4) and Leu^235^-Lys^239^ (unstructured loop helix α4-β9 strand). Taken together, this analysis provides insight into a potential stable structure rearrangement or oligomerization of W/N/L-mVP40 in presence of PS-containing vesicles that display a slow HD exchange compared to WT and previously analyzed mutants.

**Fig. 4.**
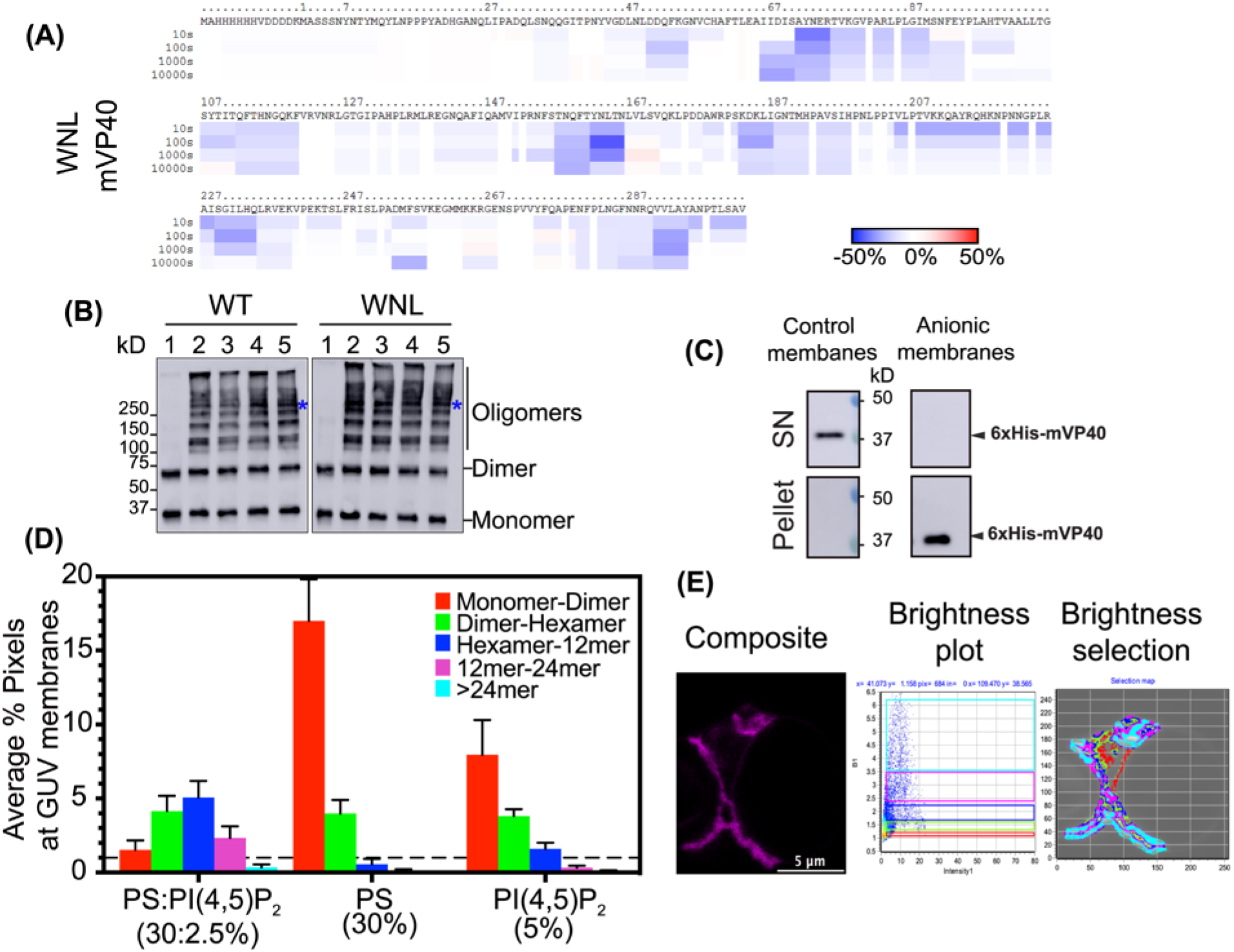
*In vitro* study of NTD/CTD oligomerization interfaces triple mutant WNL-mVP40. **(A)** Ribbon maps of W83R/N148A/L226R (WNL) mutant, indicating the difference in deuteration percentage in the presence of PC:PS (55%:45%) liposomes over the entire exchange period. Each row corresponds to each time point from 10 to 1000 seconds. Color coding: blue indicates the regions that exchange slower and red indicates the regions that exchange faster in the presence of liposomes. **(B)** *In vitro* crosslinking indicates potential oligomerization mutant still capable of higher ordered structures in the presence of anionic liposomes. Lane 1: PC (100%), lane 2 PC:PS (60:40%), lane 3: PC:PI(4,5)P_2_ (92:7.5%), lane 4: PC:PS:PI(4,5)P_2_ (75:20:5%) and lane 5: PC:PI(4,5)P_2_ (90:10%). Blue asterisk indicates a potential hexamer size of mVP40. **(C)** Liposome sedimentation assay of WNL-mVP40 was performed using control membranes (no anionic lipids) or anionic membranes (30% PS:2.5% PI(4,5)P_2_). **(D)** oligomerization profile of WNL according to different anionic membranes 30%PS:2.5%PI(4,5)P_2_ (molar ratio), 30% PS only and 5% PI(4,5)P_2_ only, determined from N&B analysis. **(E)** Representative original composite of the time-lapsed images (left panel), the number of pixels vs. intensity plot (middle panel) and brightness selection plot of the 30%PS:2.5%PI(4,5)P_2_-containing GUV (right panel).

To test this hypothesis and determine the ability of the triple mutant to oligomerize in the presence of anionic lipids, we conducted an *in vitro* oligomerization assay with the water soluble chemical crosslinker BS^3^ (Fig. 4B). WT and W/N/L proteins exhibited no oligomerization with control lipids (no anionic lipids) as expected and only the dimer and monomer were detected (Fig. 4B, lane 1). WT-mVP40 displayed different oligomers without predominance of specific molecular sizes in 40% PS containing membrane as well as 7.5% PI(4,5)P_2_ (Fig. 4B, lanes 2 and 3). However, in membranes containing PS:PI(4,5)P_2_ (20:5% mol), a band at a molecular weight >250 kDa was more obvious (Fig. 4B, lane 4, blue asterisk). The same profile was also observed in membranes containing equivalent percentage of negative charge (10% PI(4,5)P_2_ that corresponds to ~40% negative charge) with also a small increase of a band between 150 and 250 kDa (Fig. 4B, lane 5, blue asterisk). According to our estimation, these two bands may correspond to: ~206 kDa (hexamer) and ~143 kDa (tetramer), respectively. Concerning the WNL-mVP40 triple mutant, the band at ~206 kDa that may correspond to the hexameric mVP40 form was detected clearly in the 4 different anionic membrane conditions (Fig. 4B lanes 2-5, blue asterisk). This data suggests that WNL-mVP40 is most likely forming a new and unique structure in presence of anionic membranes as a result of the three mutations. Furthermore, WNL-mVP40 had a similar ability to bind PS-PI(4,5)P_2_ membranes compared to WT-mVP40, as shown in the liposome sedimentation assay (Fig. 4C, 4D). As expected, WNL-mVP40 did not bind control membranes (neutral) but showed a normal binding to anionic membranes indicating that the triple mutation did not affect the lipid binding efficiency of the protein.

To assess the abundance of particular oligomers of WNL-mVP40 in the presence of anionic membranes, we performed *in vitro* N&B analysis with GUVs as described above. The data is summarized in Fig. 4D and Table S3 with respect to the oligomerization profile of WNL-mVP40 in the presence of different negatively charged membranes while Fig. 4E shows the ability of the mutant to bind and enrich efficiently at PS:PI(4,5)P_2_ membranes (composite panel). In these analyses, we demonstrated that ~13% of total pixels detected were enriched protein at the membrane of the GUVs containing either PS:PI(4,5)P_2_ or PI(4,5)P_2_ only. However, the oligomerization profiles of the protein were different in the two lipid membranes. In short, PS:PI(4,5)P_2_ bound protein formed mostly hexamer-12mer (~37.8% of total bound protein), 30.88% dimer-hexamer, 17.4% 12-24mer, 11.36% monomer-dimer and about 3.4% were larger oligomers (>24mer, Fig. 4D, 5F). On the other hand, on PI(4,5)P_2_-containing membranes, WNL-mVP40 was mostly abundant as monomer-dimer with about 57.7% of total bound protein, ~27.66% dimer-hexamer, 11.57% hexamer-12mer, ~2.4% 12mer-24mer and a very small population could be detected as >24mer (less than 1%). Furthermore, the same analysis was performed with PS-containing membranes, and as expected, the mutant displayed mostly a monomer-dimer profile at the GUV membrane with more than 79% of total bound protein, 18% were dimer-hexamer and 2.3% were hexamer-12mer (Fig. 4D). All together, these analyses indicate that WNL-mVP40 exhibited an oligomerization profile comparable but more exaggerated to the NTD oligomerization interface mutant W83R/N148A-mVP40 (Fig. 3E; 4D). Altogether, *in vitro* analysis of NTD/CTD oligomerization interfaces demonstrate they are not involved in the ability of mVP40 to bind anionic membranes, however, both the NTD and CTD interfaces are required for efficient protein oligomerization at the membrane and matrix assembly.

**Fig. 5.**
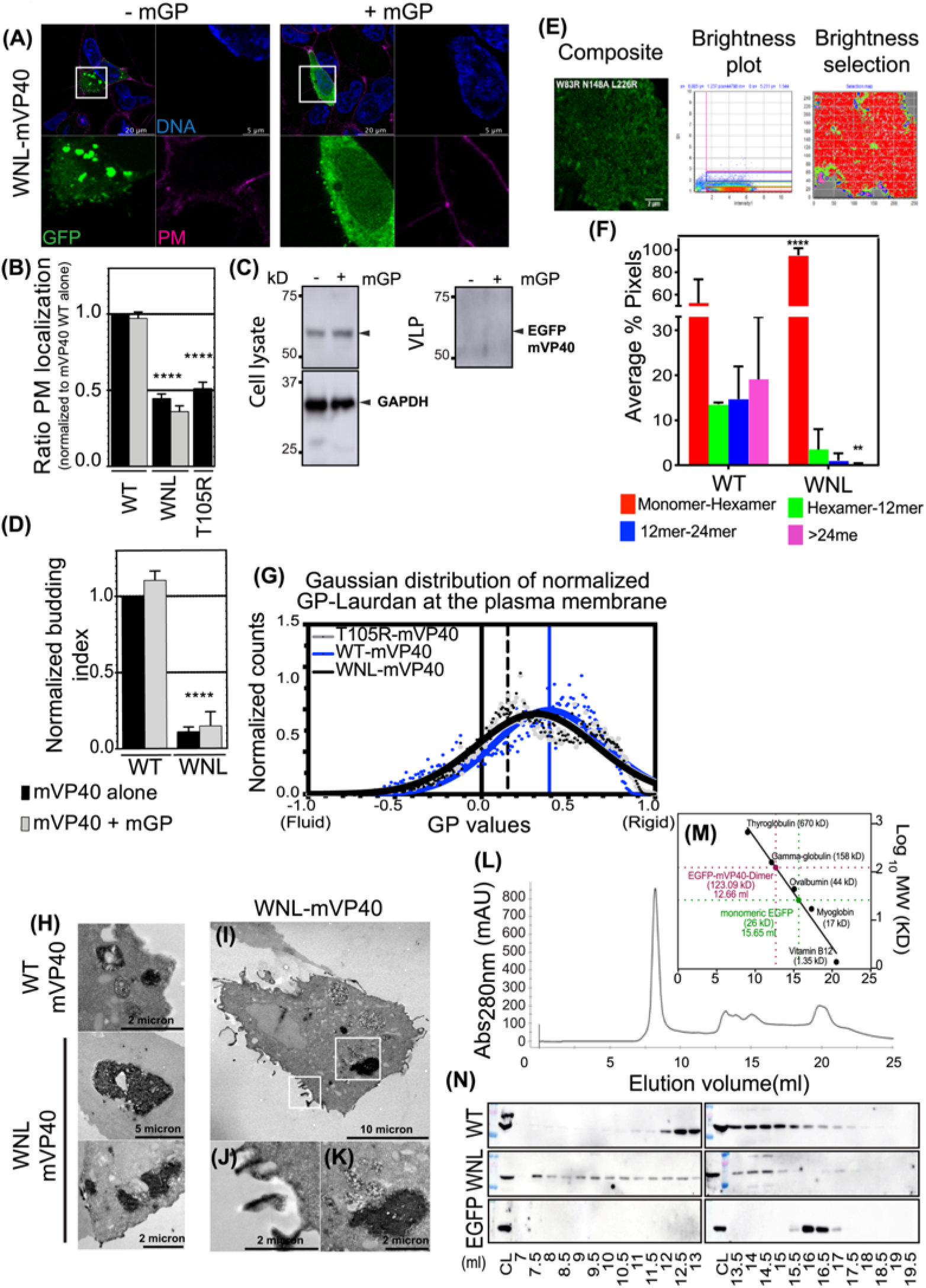
Cellular behavior of WNL-mVP40 mutant. **(A)** HEK293 cells, expressing EGFP-constructs +/-mGP, stained for DNA (blue) and PM (pink). **(B)**Ratios of PM retention represented as averages ± S.E.M of three independent means. WT-mVP40 data are extracted from Fig. 1B. Statistical analysis was performed as described in Fig 2 (\*\*\*\**p*<0.0001). **(C)** Western blot assay performed on cells and VLP quantified in **(D)** as described in Fig. 3. **(E)** N&B analysis of cellular EGFP-WNL-mVP40 24 h.p.t. **(F)** Average % pixels of estimated oligomerization forms of EGFP-WT and WNL-mVP40. **(G)** Gaussian fitted normalized GP distribution curves of laurdan across PM of cells expressing EGFP-WNL-mVP40 (black) compared to WT (blue) and T105R-mVP40 (grey) as described in Fig 2. TEM micrographs of cells co-expressing GBP-APEX2 and EGFP-mVP40 (WT or WNL) in **(H)**, while **(I)** and insets **(J)** and **(K)** show the structure of intracellular WNL protein aggregations. **(L)** The chromatogram of gel filtration analysis of protein extract from HEK293 cells transfected with EGFP-WT-mVP40 shown as absorbance (280 nm) versus elution volume. Molecular mass standard curve is plotted in **(M)** as log values of molecular weights versus elution volume. **(N)** Western blot analyses of each protein are indicated. EGFP empty vector served as a negative control. CL: cell lysate.

### NTD/CTD oligomerization interfaces triple mutant is unable to localize and oligomerize at the plasma membrane

Next, we generated an EGFP tagged triple mutant of both NTD and CTD oligomerization interfaces (EGFP-WNL-mVP40) to expand on the involvement of these two interfaces in plasma membrane localization. As shown in Fig. 5A, the triple mutant was unable to localize to the plasma membrane similarly to the monomeric mutant T105R (Fig. 2A), which was corroborated by the quantitative analysis (Fig. 5B). Furthermore, the VLP budding efficiency was tested by functional budding assays and indicated that W/N/L-mVP40 was unable to bud from the plasma membrane (Fig. 5C, 5D). In both assays, co-expressing the mutant with mGP did not rescue the WT phenotype (Fig. 5A bottom panel, Fig. 5C, 5D) indicating that the trafficking and stabilization at the plasma interface was highly dependent on the oligomerization efficiency of mVP40. To confirm that WNL-mVP40 is unable to oligomerize at the plasma membrane, we performed N&B analysis in living cells. Figure 5E and 5F revealed that oligomerization of WNL was abrogated, where hexamer-12mer represented only 3.52%, 12mer-24mer 1.05% and >24mer 0.21% of total pixels detected. Likewise, a significant increase in monomer-hexamer was observed with up to 95% of the total pixels (Fig. 5F, Table S2). Taken together, this analysis supports the requirement of both NTD and CTD oligomerization interfaces for the correct and efficient binding of mVP40 to the plasma membrane of host cells and productive homo-oligomerization to form the viral matrix needed for VLP budding. Based on this phenotype, it was of our interest to know if W/N/L mutations had an effect on plasma membrane fluidity. Laurdan imaging analysis described in Fig. 5G (images in Fig. S4) highlighted the ability of WNL-mVP40 to induce a mild increase of rigidity at the plasma membrane, albeit slightly less than WT-mVP40. Interestingly, the phenotype was identical to the monomer mutant T105R-mVP40 with a GP index about 0.3. These data suggest that cellular expression of mVP40 may affect the plasma membrane lipid composition or distribution even before mVP40 resides there.

During our analysis of WNL-mVP40, we observed intracellular vesicular structures within transfected cells and rarely, the protein able to reach the plasma membrane (data not shown). To investigate these observations further, TEM analysis on cells co-expressing EGFP-WNL-VP40 with GBP-APEX2 was performed. Figure 5H-K is a representative micrograph of cells co-expressing W/N/L-mVP40 and GBP-APEX2. Trace levels of APEX2 signal were detected at the cell periphery (Fig. 5H, 5J). However, a large accumulation of APEX2 signal was observed in the cytosol (Fig. 5H, 5I, 5K). A similar accumulation was observed previously for WT-mVP40 (Koehler *et al.*, 2018) in addition to our TEM experiments (Fig. 5H) and confocal imaging (data not shown). In Figure 5H, we compared the structure of the intracellular accumulations in both WT (top panel) and WNL-mVP40 (middle and bottom panels). We noted that these protein accumulations were more abundant, larger and less structured in the triple mutant compared to WT.

At this point it was necessary to examine whether the triple mutant displayed a specific oligomerization profile in cells and not only at the plasma membrane. For this purpose, we performed a size exclusion (SEC) assay on protein extract from cells expressing either EGFP-WT-mVP40, or EGFP-WNL-mVP40 as described previously (Liu *et al.*, 2010). Monomeric EGFP alone was used as a control of a protein that does not oligomerize. In brief, cells transiently expressed different constructs for 24 hours and proteins were extracted in 1% triton X-100 prior to separation by SEC (Fig. 5L-N). Internal molecular weight standards were also used for molecular weight estimation (Fig. 5M). EGFP-WT-mVP40 was detected in different fractions that correspond to the peaks at elution volumes 12.5 to 13.5 ml and 14 to 15 ml (Fig. 5N). These peaks likely correspond to dimeric and monomeric forms, respectively. In addition, a very small amount of protein was detected at elution volumes 10.5 to 11.5 ml that should correspond to larger oligomers, most likely hexamers. In contrast, EGFP-WNL-mVP40 was detected predominantly in fractions at elution volumes 8.5 to 10 ml along with elution volumes from 12.5 to 13.5 ml and 14 to 15 ml, corresponding most likely to larger homo-oligomeric, dimeric and monomeric forms (Fig. 5N). Interestingly, the protein was also detected in the void volume (elution volume 7 ml), which may indicate the presence of very large oligomers or aggregates (Fig. 5N). These data suggest that the NTD/CTD oligomerization interface triple mutant in the cell is forming the dimer, but also larger oligomers that block its trafficking to the plasma membrane. EGFP extract eluted from the FPLC column in one peak at elution volume 15.5 to 16.5 ml, that correspond to monomeric form as expected, supporting that the previous observations are a result of the homo-oligomerization of mVP40.

### MD simulation of NTD and CTD oligomerization interfaces

To characterize the differences in the oligomer interfaces of eVP40 and mVP40 (Fig. 6A-B), we calculated the distance between the tryptophan residues (Trp^95^-Trp^95^ for eVP40 and Trp^83^-Trp^83^ for mVP40) at the interface. Upon relaxation with MD simulation, initially separated Trp^83^ residues in the modeled mVP40 oligomer interface get closer and interact with each other (Fig. 6A, right panel). This is shown by the significant decrease in the Trp^83^-Trp^83^ distance, whereas the W95 residues in eVP40 remain separated and the Trp^95^-Trp^95^ distance does not change during the simulation window (Fig. 6C). In addition, Asn^148^ is in close proximity to make occasional backbone hydrogen-bonding with Ile^88^ at the interface.

**Fig. 6.**
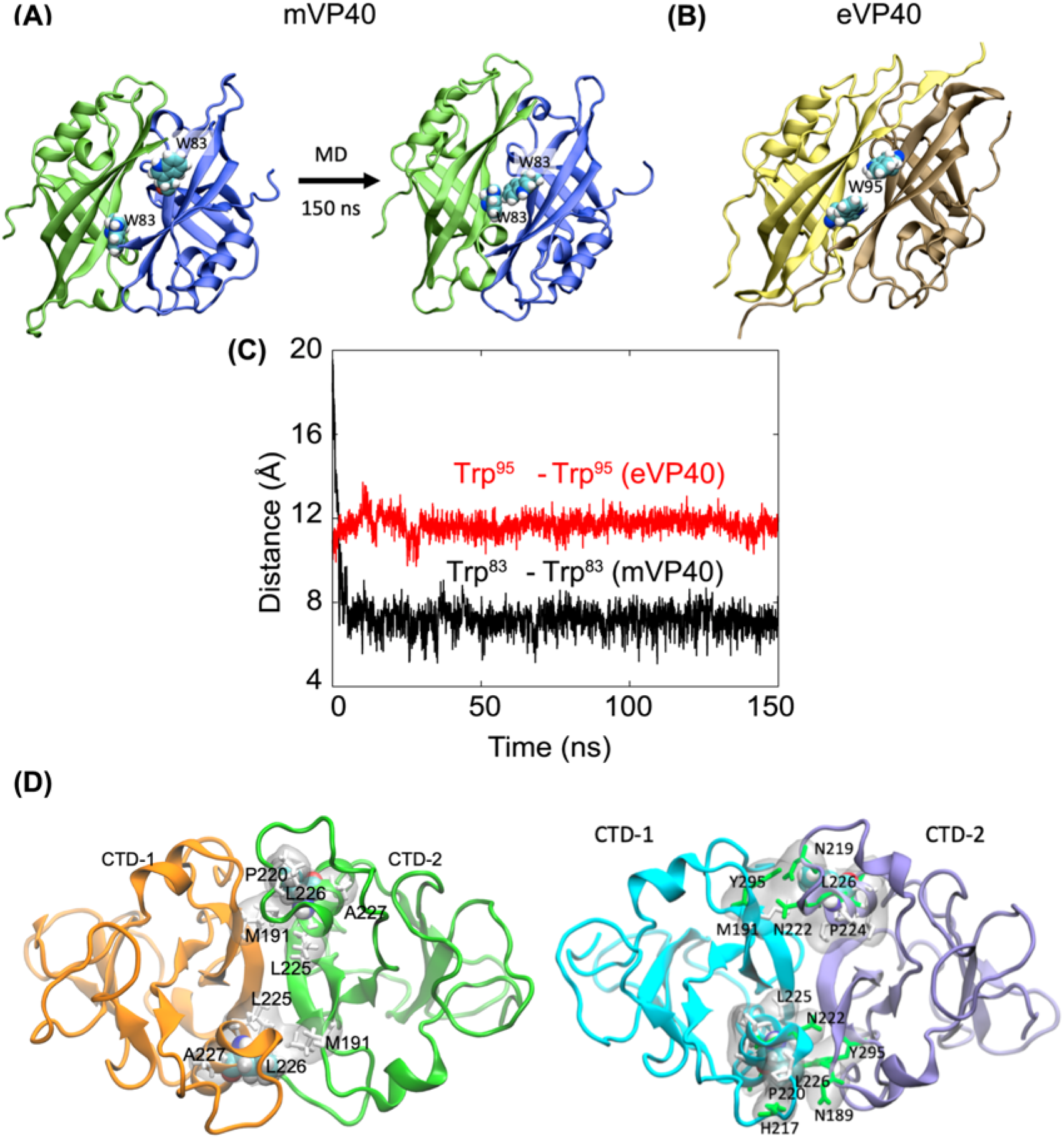
Molecular dynamics simulations of the oligomer interfaces of the mVP40. **(A)** The mVP40 oligomer interface modeled based on eVP40 structure initially shows separated W83 residues as in eVP40 (Trp^95^) shown in **(B)**. However, upon 150 ns MD simulation, the structure relaxes so that the interface residues W83 interact with each other. **(C)** Center of mass distance between Trp^83^ residues in mVP40 (black curve) and between W95 residues in eVP40 (red curve) as a function of time. **(D)** Hexamer-hexamer interface in the mVP40 filament (CTD from each monomer is in showed in different colors). The hydrophobic residues within 3 Å of Leu^226^ at the mVP40 hexamer-hexamer interface are highlighted. The hydrophobic interaction at the hexamer-hexamer interface may provide an agile interface, giving flexibility to the filaments. **(E)** Zoom into hexamer-hexamer interface in the mVP40 filament formed through CTD-CTD linear oligomerization as proposed by Wan *et al.* (Wan *et al.*, 2020). (CTD from each monomer is in showed in different colors)

To investigate the interactions at the CTD-CTD interface formed between the two hexamers shown in Fig. 1A, we simulated the CTD-CTD complex (Fig. 6D) for 100 ns. Similar to eVP40, this interface consists of primarily hydrophobic residues including Leu^226^, Pro^220^, Met^191^, Ala^227^, and Leu^225^. Therefore, Leu^226^ is part of the hydrophobic core at the interface that provides stability as well as flexibility to the CTD hexamer-hexamer interface.

## Discussion

mVP40 is described as an anionic charge sensor with lack of stereospecificity to PI(4,5)P_2_ at the plasma membrane compared to eVP40, which requires PI(4,5)P_2_ for proper binding and matrix assembly (Johnson *et al.*, 2016; Wijesinghe and Stahelin, 2016). However, it is still not known how the oligomerization of mVP40 occurs to undergo matrix assembly and what role anionic lipids play in promoting the proper mVP40-mVP40 oligomerization during the virus assembly. HDX-MS analysis previously conducted on mV40 in the presence and absence of anionic lipids revealed two potential oligomerization interfaces (Wijesinghe *et al.*, 2017). The NTD oligomerization interface was proposed to include β2, β3 (Trp^83^ residue), and β7 antiparallel β sheet structures and the CTD oligomerization interface was proposed to include the α4 helix (Leu^226^ residue) similarly to higher-ordered oligomerization of eVP40 hexamers (*via* CTD end-to-end contacts as previously described (Bornholdt *et al.*, 2013)). These two regions of mVP40 exhibited a reduced deuteration level in presence of anionic lipids (Wijesinghe *et al.*, 2017). Furthermore, both NTD and the CTD oligomerization interfaces are hydrophobic suggesting multimerization driven by hydrophobic interactions. It is also possible that each interface is involved in a specific lipid-dependent oligomerization pattern of mVP40. To better understand the mechanism of these potential hydrophobic interactions at NTD and CTD, we replaced the residues Trp^83^ in NTD and Leu^226^ in CTD with the charged amino arginine to repulse protein-protein interactions in these regions.

In this study, the cellular analysis of NTD oligomerization interface double mutant W83R/N148A indicated inability to enrich at the plasma membrane compared to WT-mVP40. Further, the double mutant had reduced higher-ordered oligomerization, significant increase of small oligomers (monomer-hexamer) and decreased budding deficiency. Similar results had been described previously on untagged or HA tagged mVP40 (Oda *et al.*, 2016; Koehler *et al.*, 2018). This mutant had been described to be able to dimerize in solution (Oda *et al.*, 2016) but no data were available on the oligomerization pattern of W83R/N148A at lipid membranes. Here, we demonstrated that the mutant is still able to multimerize (hexamer-12mer) but deficient to form higher-ordered oligomers at the plasma membrane of transfected cells. This inability of the NTD mutant can explain the deficiency in VLP formation and its low enrichment at the plasma membrane. This was also observed *in vitro* using PS:PI(4,5)P_2_-containing GUVs where only a small population (3.5 fold less than WT) could enrich at the vesicle’s membranes suggesting proper enrichment and assembly at these membranes requires proper mVP40 NTD-mediated oligomerization. The single mutant W83R showed a similar oligomerization profile to WT but still was unable to form VLPs. This may suggest compensation in the Trp^83^ mutant by the adjacent residue (Asn^148^), however, the NTD-NTD oligomerization through Trp^83^ is a key process for VLP elongation and release. Moreover, the mutations at the NTD oligomerization interface had a comparable effect on the membrane rigidity increase compared to WT-mVP40 probably due to lipid rearrangement and/or clustering at the plasma membrane upon protein oligomerization. Lipid rearrangement and clustering (e.g., domain formation) is often required for virus particle budding (Krementsov *et al.*, 2010; Hogue *et al.*, 2011; Mücksch *et al.*, 2017; Madsen *et al.*, 2018) and a similar phenomena has been proposed at the inner leaflet of the plasma membrane where eVP40 hexamer significantly enhanced PI(4,5)P_2_ clustering (GC *et al.*, 2016) In this present study, we can’t totally omit the ability of NTD oligomerization interface mutant to form proper VP40 hexamer structure but our data clearly demonstrate a significant deficiency in forming a functional viral matrix.

The CTD region of mVP40 contains two basic loops (1 and 2) involved in anionic lipid interactions. Previous HDX-MS studies highlighted the potential involvement of α4 helix at the CTD region, including residues Leu^226^ and potentially Ser^229^ in hexamer-hexamer interactions (Wijesinghe *et al.*, 2017). The impacts of mutations on the CTD region were non-significant (L226R) or mild (L226R/S229A) on cellular localization, protein oligomerization and virus-like particle release. Importantly, in presence of the MARV glycoprotein, L226R mutant had reduced plasma membrane localization compared to WT-mVP40 and consistently this resulted in a reduction in VLP production. Furthermore, membrane fluidity analysis demonstrated that both L226R and L226R/S229A are unable to induce changes in plasma membrane rigidity compared to EGFP controls. Thus, we hypothesize that CTD-CTD oligomerization of mVP40 is important but not required to stabilize the matrix assembly that can result in lipid rearrangements at the plasma membrane. Our *in vitro* oligomerization assay of L226R mutant supports this hypothesis. Indeed, L226R enriched 1.5 fold less than the WT-mVP40 at PS:PI(4,5)P_2_ containing membranes and more than 50% of enriched protein were small oligomers (dimer to hexamer) with a consistent decrease of more ordered oligomers (hexamer-12mer, 12mer-24mer and >24mer). Thus, larger mVP40 oligomers, which are attributed to CTD-CTD are more likely to alter the plasma membrane rigidity.

Investigating the role of each phospholipid in the *in vitro* oligomerization of WT-mVP40 and the mutants was in our opinion critical to better understand the involvement of each oligomerization interface in MARV matrix assembly. This simplified system with GUVs containing only PS, only PI(4,5)P_2_ or both PS and PI(4,5)P_2_, provided an important understanding of anionic lipid-dependent mVP40 oligomerization. First, in PS-containing GUVs, mVP40 forms mostly small oligomers (from monomer to hexamer) with a very small population (9.4%) of hexamer-12mer. This clearly demonstrates that upon binding to PS, mVP40 clusters at the membrane without further high-ordered oligomerization suggesting that NTD-NTD oligomerization is more prominent in presence of PS alone. Next and in PI(4,5)P_2_-containing GUVs, mVP40 was able to form higher-ordered oligomers up to 24mer compared to the previous conditions with a decrease of dimer-hexamer population. This indicated that upon PI(4,5)P_2_ binding, mVP40 may undergo conformational changes that promote CTD-CTD interaction for high-ordered oligomerization. If our predictions on the role of Trp^83^ and Asn^148^ in mediating NTD-NTD oligomerization and of L226 in CTD-CTD oligomerization, the W83R/N148A mutant is most likely to exhibit a deficient oligomerization profile in PS-containing membranes, while L226R should have oligomerization defects in PI(4,5)P_2_-containing membranes. The relative decrease of oligomerization of W83R/N148A was indeed observed in PS-containing GUVs with a significant accumulation of monomeric-dimeric protein at the membrane. However, L226R showed a very pronounced increase of dimer-hexamer population in PS:PI(4,5)P_2_ but not in PI(4,5)P_2_-containing membranes. This suggested that the role of CTD-CTD interactions is more important in membranes close to physiological compositions and helps explain the inability of L226R and L226R/S229A mutations to increase plasma membrane fluidity upon mVP40 binding and assembly.

In the present study, we provided insight on the potential role of NTD and CTD interfaces in mVP40 membrane enrichment, protein oligomerization and matrix assembly, and VLP budding. Our *in vitro* analyses using anionic lipid-containing vesicles highlighted the structural changes that CTD and NTD oligomerization interfaces undergo upon lipid binding and oligomerization. It is not completely clear to us how and when the CTD-CTD oligomerization occurs. CTD-CTD-interactions may be required at a specific stage of the matrix assembly after protein associates with the plasma membrane and establishment of NTD-NTD interactions to initiate the protein high ordered-oligomerization. A recent model proposed CTD-CTD linear oligomerization that is most likely in both MARV and EBOV virions and VLPs (Wan *et al.*, 2020). However, our study suggests the importance of NTD-NTD oligomerization in cells and *in vitro* to establish the building blocks for higher-ordered oligomer formation and particle release. It is possible that NTD-NTD interactions are required to increase membrane bending, elongation of tubule and/or for host cell factor recruitment at assembly sites. Furthermore, using the recent model proposed in CTD-CTD linear oligomerization ((Wan *et al.*, 2020), Fig. S6), we simulated the CTD-CTD complex that indicated this interface may involve Met^191^, Asn^222^, Tyr^195^ and Leu^226^ as we report here (Fig. 6E). Future studies aimed at mutations of this region should help to clarify the detailed interactions that talk place at these CTD-CTD interaction sites. Finally, the triple mutant WNL-mVP40 showed a completely different phenotype and behavior in cells or *in vitro*. The ability of the triple mutant to bind lipids efficiently and oligomerize suggests an uncommon and uncharacterized homo-oligomerization in cells and with lipid membranes, involving non-studied residues, that seemingly blocks the protein trafficking to the plasma membrane. Further analysis of structural rearrangement of this mutant can provide precious information on potential oligomerization of mVP40 required for cell signaling and/or trafficking.

Taken together, this study demonstrated that mVP40 has two oligomerization interfaces at NTD and at CTD. Each interface regulates specific protein oligomerization at the plasma membrane in a lipid-dependent manner, membrane fluidity changes, matrix assembly, VLP elongation and budding. Thus, small molecule or other therapeutic agents can be considered to disrupt the inter and intramolecular interactions of mVP40 to block the proper viral matrix assembly and prevent release of virus progeny.

## Materials and Methods

### Site directed mutagenesis

Site directed mutagenesis was performed using a Q5® Site-Directed Mutagenesis Kit (New England Bio labs) using primers listed in Table S1 according to the manufacturer’s protocol. The same primer sets were used to generate mutants with pcDNA3.1-EGFP-WT-mVP40 or a His_6_-tag or EGFP tag in pET46 with the His_6_-WT-mVP40 vector originally a kind gift from Dr. E. Ollmann Saphire (La Jolla Institute for Immunology).

### Cell culture and live cell imaging

COS-7 or HEK293 cells were maintained in DMEM (Corning, NY) containing 10% FBS and 1% Penicillin/streptomycin at 37°C in a 5% CO_2_ humidified incubator. Cells were grown until 70% confluency before transfection in 8 well Nunc Lab Tek II chambered slides with 0.16 mm cover glass thickness from Thermo Fisher Scientific (Waltham, MA). Transfections were performed using Lipofectamine 2000 or Lipofectamine LTX and PLUS reagents (supplied ThermoFisher Scientific) according to the manufacturer’s protocol.

The enhanced green fluorescent protein (EGFP) signal was imaged 14 hours post transfection (performed at 37°C) on a Nikon Eclipse Ti Confocal inverted microscope (Nikon, Japan), using a Plan Apochromat 60x 1.4 numerical aperture oil objective or a 100x 1.45 numerical aperture oil objective, respectively. Cells were stained for 15 min at 37°C with 5 μg/ml Hoechst 3342 and 5 μg/ml wheat germ agglutinin, Alexa Fluor 647 conjugate (WGA-Alexa Fluor 647, Molecular Probes^TM^) in growth media, for nucleus and plasma membrane staining, respectively. Cells were imaged using the 405 nm, 488 nm and 647 nm argon lasers to excite Hoechst, EGFP and WGA-Alexa Fluor 647, respectively. Plasma membrane localization ratios were calculated using the integrative density intensities at the plasma membrane determined using the WGA-Alexa Fluor 647 signal compared to the total intensities of the entire cell using ImageJ(Rasband, 2015).

### Functional budding assays and Western blotting

Functional budding assays were adapted from an established protocol (Harty, no date). HEK293 cells at 1-1.5 × 10^6^ density, were transfected with EGFP-mVP40 constructs with or without co-expression of mGP using Lipofectamine LTX and PLUS reagent according to the manufacturer’s protocol. At 24 hours post transfection, the media containing virus-like particles (VLP) were harvested and collected as previously described (Oda *et al.*, 2016). Total protein contents (5 μg) from cell lysates and VLP samples were resolved on a 12% SDS-PAGE gel (protein amount appropriate for 15 well gels) prior to transferring on nitrocellulose membrane. Target proteins were detected using indicated primary antibody, 1:200,000 dilution of Rabbit α-mVP40 (IBT BioServices) and in some experiments 1:2000 dilution of Mouse α-GFP (ThermoFisher), Mouse α-GAPDH (ThermoFisher) was used at 1:10,000 final dilution, followed by the appropriate secondary antibodies horseradish peroxidase (HRP) conjugated, Goat α-Rabbit or Sheep α-Mouse (Abcam) at 1:5,000 final dilution for both. HRP signal was detected using Amersham Prime ECL reagent (GE Lifesciences, Chicago, IL) and imaged on a Amersham Imager 600. VLP budding index of different mVP40 proteins was performed with densitometry analysis using ImageJ (Rasband, 2015). The following equation was applied:

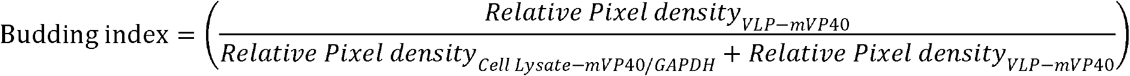

The budding index of each mutant was normalized to the WT-mVP40 budding index.

### Transmission electron microscopy: Chemical fixation and APEX processing

5.2 ×10^5^ of HEK293 cells were seeded on 25 mm diameter poly-L-lysine coated cover glass. The next day, 2.5 μg of each APEX2-csGBP plasmid and mVP40 constructs were co-transfected using Lipofectamine LTX reagents cells were incubated at 37°C, 5% CO_2_ for 6 hours, then the transfection medium was changed for DMEM (Corning, NY) containing 10% FBS. Cells were then incubated at 37°C, 5% CO_2_ for 8 hours, after which time cells were rinsed with Dulbecco’s 1x PBS and were chemically fixed with 2.5% glutaraldehyde in 0.1M cacodylate buffer for 30 min. Fixed cells were then rinsed 3x for 5 min each with 0.1 M cacodylate buffer and washed with 1 mg/mL of 3,3’-diaminobenzidine (DAB) (Sigma-Aldrich) in cacodylate buffer for 2 min. Following the wash, cells were incubated in a freshly made solution of 1 mg/mL of DAB and 5.88 mM of hydrogen peroxide in cacodylate buffer for 25 min on ice. Cells were washed 3x for 5 min each with cacodylate buffer, incubated in an aqueous solution of 1% osmium tetroxide for 10 min and then washed with distilled water. Dehydration was conducted using increasing concentrations of ethanol (50%, 75%, 95%, and 100% made from 200 proof ethanol), transitioned using 100% acetonitrile and followed by resin infiltration of the cells using increasing concentrations of Embed 812 Epoxy resin without the accelerator in acetonitrile (2:1 and then 1:2), and finally with Embed 812 containing the accelerator. Coverslips were then embedded on resin filled beam capsules (cell-face-down) and incubated in an oven at 60◻ for 24 hrs. After polymerization, coverslips were removed by dipping the coverslip faced block in liquid nitrogen. Serial sections were then collected by sectioning the block samples en face and ribbons were collected on formvar-coated slot grids.

Thin (90 nm) serial sections were obtained using a UC7 ultramicrotome (Leica) and collected onto formvar-coated copper slot grids (EMS). Glass knives were prepared for trimming, while an Ultra 35° diamond knife (Diatome) was used for sectioning the block samples. Sections were screened on a Tecnai T-12 80kV transmission electron microscope and average 10-15 cells were visualized from each sample.

### Number & Brightness (N&B) analysis on mammalian cells

Number & Brightness (N&B) experiments were performed as described previously (Adu-Gyamfi *et al.*, 2012a; Johnson *et al.*, 2016; Bobone *et al.*, 2017). HEK293 cells were seeded onto 1.5 mm poly-D-lysine coated coverslips with 0.17 mm thickness in 6-well plates at 70% confluency. Cells were transfected with either EGFP or EGFP-tagged mVP40 constructs as described previously. Cell were washed 24 hours post transfection with 1x PBS, transferred to an Attofluor^TM^ chamber (Invitrogen), and imaged in Live Cell Imaging Solution (Gibco, Life Technologies, Carlsbad, CA) using the Zeiss LSM 880 upright microscope (Carl Zeiss AG, Germany) and a LD “C-Apochromat” 40x/1.1 W Corr M27 objective and a 488◻nm argon laser to excite EGFP. Each image was acquired using the same laser power (0.01), resolution (256×256), pixel dwell time (16 μs), frames (50), and zoom (pixel size of 50 nm). SimFCS Globals Software (Laboratory for Fluorescence Dynamics, University of California, Irvine, CA) was used for analysis.

On each experimental day, EGFP expressing cells were imaged and SimFCS4 software (G-SOFT Inc.) was used to determine the true brightness (B) of monomeric EGFP (0.058-0.13), which is consistent with previous analyses (Youker and Teng, 2014). To calculate the apparent brightness value of mVP40 oligomers, the B_monomer_ value was multiplied by the corresponding oligomer value (i.e. dimer = 2, hexamer = 6). Using SimFCS, bins were placed in the brightness plot to correspond with the respective oligomer size. The number of pixels of monomer-hexamer, hexamer-12mer, 12mer-24mer, and 24mer+ bins were recorded. Average % pixels of each oligomeric state was ratiometrically determined by the total number of pixels in each bin vs. the total number of pixels in the image.

### Laurdan and membrane fluidity analysis

Membrane fluidity analysis was performed according to Owen *et al.* (2012) (Owen *et al.*, 2012). In brief, 14 hours post transfection of HEK293 cells with different mVP40 constructs or EGFP plasmid, cells were treated with 10 μM laurdan (Invitrogen^TM^, stock made in DMSO at final concentration of 5 mM) in culture media and incubated for 30 min at 37°C in a humidified 5% CO_2_ atmosphere. Cells were then imaged with a Ti-E inverted microscope equipped with Nikon’s A1R confocal and a Spectra Physics IR laser tunable to 800 nm for multi-photon confocal imaging of the laurdan dye and images collected with photon multiplier tubes (PMT) set at 400–460 nm and 470–530 nm for ordered (PMT1) and disordered (PMT2) membranes, respectively. Calibration images were acquired with 100 μM laurdan in culture media to calculate the measured generalized polarization factor (GP). Image processing was done using ImageJ and GP distribution was determined using the Laurdan_GP macro provided in (Owen *et al.*, 2012).

### Gel filtration analysis of EGFP and EGFP tagged WT and mutant mVP40 protein

Human HEK293 cells were transfected with EGFP constructs 24 hours prior to protein extraction described previously in Liu et al. (Liu *et al.*, 2010). In brief, cells were washed with PBS and lysed with PBS containing 1% triton X-100, scrapped, collected and incubated on ice for 10 min. Lysates were cleared by centrifugation at 2,000 rpm for 10min at 4°C and filtrated through a 0.22-μm-pore-size filter. The cleared protein extract was then separated according to protein sizes on Superdex^TM^ 200 Increase 10/300 GL, fast-protein liquid chromatography (FPLC) column using ÄKTA pure (GE healthcare). Eluted proteins were collected in 0.5-ml fractions and analyzed by SDS-PAGE and then by Western blotting with anti-EGFP antibody, as described above. The chromatogram plotting absorbance (280 nm) versus elution volume was generated with Unicorn 7.2 software.

### Protein Purification

Purification of mVP40 wild type, mutants (W83R/N148A, L226R, W83R/N148A/L226R) and His_6_-tag alone proteins was adapted from a previously established protocol (Wijesinghe and Stahelin, 2016). In brief, protein expression was performed over night at 18°C with 250 μM IPTG at an optical density (OD_600nm_) from 0.7 to 0.8. The bacteria pellets were lysed for 30 min on ice in lysis buffer: 20 mM Tris pH 8.0, 500 mM NaCl, 1x halt protease inhibitors, 300 μg/ml lysozyme, 100 μg/ml RNAse and 3 μM phenylmethylsulfonylfluoride (PMSF, Thermo Fisher Scientific, Waltham, MA). The lysis solutions were then subjected to 5 sonication cycles at 38% (10 sec ON, 59 sec OFF). After 1 hour centrifugation at 15,000 x *g* at 4°C to clarify the lysate from cell debris and membranes, the protein solutions were incubated with Ni-NTA agarose for 30 min at 4°C with continuous rocking. The proteins were washed with 20 mM Tris pH 8.0, containing 500 mM NaCl and 50 mM imidazole prior to three 5 min stepwise elutions with 20 mM Tris pH 8.0, containing 500 mM NaCl and 300 mM imidazole. The mVP40 eluted fraction were washed and dialyzed against storage buffer 20 mM Tris pH 8.0, containing 500 mM NaCl and 20 % glycerol using 30K MWCO concentration tubes (or 3K MWCO for His-tag alone purification). The protein purity and enrichment were confirmed by SDS-PAGE and size exclusion using a HiLoad® 16/600 Superdex® 200 pg column using an ÄKTA pure (GE healthcare). However, for *in vitro* assays with lipids, the proteins were used post dialysis.

### Liposome Sedimentation Assays

All lipid used here were purchased from Avanti Polar Lipids, Inc. (Alabaster, AL). Large unilamellar vesicles (LUV) were used for liposome sedimentation assays. Lipid mixtures were prepared at the indicated compositions and chloroform soluble lipids were dried to form lipid films under a continuous stream of N_2_. In each experiment, addition of anionic lipids was compensated with an equal mol% decrease in POPC, while POPE (9%) and dansylPE (1%) were held constant. Lipid films were then hydrated in liposome sedimentation buffer (260 μM raffinose pentahydrate in PBS, pH 7.4), vortexed vigorously, and extruded through a 200 nm Whatman polycarbonate filter (GE Healthcare) after incubation at 37°C. Vesicle size was confirmed by dynamic light scattering using a DelsaNano S Particle Analyzer (Beckman Coulter, Brea, CA). LUV solutions were diluted 4 times in PBS (pH 7.4) to reduce the raffinose pentahydrate concentration, and LUVs were pelleted at 50,000 x *g* (22°C) for 15 min. The supernatant was discarded and the raffinose filled-LUVs were resuspended in PBS (pH 7.4).

Liposome sedimentation assays were performed as described previously (Julkowska, Rankenberg and Testerink, 2013). In brief, protein and LUVs were mixed for final concentrations of 5 μg/ml and 2 mM respectively, and incubated for 30 min on ice. Following incubation, protein bound-LUVs were pelleted (16,000 x *g*, 4°C, 30 min), and the supernatants containing unbound proteins were transferred into new tubes. The protein bound-LUV pellet was washed in PBS and pelleted again (16,000 x *g*, 4°C, 30 min). The supernatant was discarded, and the pellet was resuspended in an equal volume as the unbound protein supernatant sample. Equal volumes of supernatant and pellet samples were resolved on a 10% SDS-PAGE gel and transferred onto a nitrocellulose membrane. The proteins were detected using the primary antibody (Mouse α-His at 1:2500 dilution, Sigma Aldrich) followed by the HRP conjugated secondary antibody (Sheep α-Mouse at 1:7000 dilution) The HRP signals were detected and analyzed as described above. To calculate %protein bound the following equation was used:

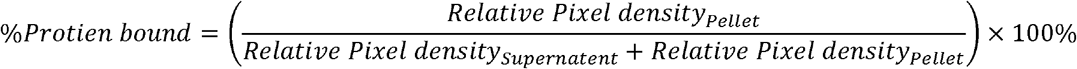

### Giant unilamellar vesicle (GUV) preparation

GUVs were prepared by a gentle hydration method (Reeves and Dowben, 1969; Darszon *et al.*, 1980; Yamashita *et al.*, 2002). Briefly, 1 mM lipid of lipid control mixture was made and contained POPC:POPE:POPS:Biotin-PC:fluorescent PC (TopFluor PC) at 59:10:30:1:0.2% molar ratio, or with 2.5% molar ratio brain phosphatidylinositol 4,5-bisphosphate PI(4,5)P_2_ added with the ratio of POPC were adjusted accordingly. PI(4,5)P_2_-containing lipid mixtures were made by mixing POPC, POPE, PI(4,5)P_2_, Biotin-PC and TopFluor PC at 84:10:5:1:0.2% molar ratio The lipid mixtures were made into a 5 mL round-bottom glass flask and the chloroform was removed with rotary movements under a continuous stream of N_2_. The lipid films were then hydrated overnight at 37°C in appropriate volume of GUV hydration buffer (10 mM HEPES, pH 7.4 containing 150 mM NaCl, and 0.5 M sucrose).

### N&B analysis on GUVs

Freshly made GUVs were diluted 10 times in GUV dilution buffer (10 mM HEPES, pH 7.4 containing 150 mM NaCl, and 0.5 M glucose) and placed on 6 mm diameter chambers made from a silicon sheet using a core sampling tool (EMS # 69039-60). The silicon chamber was mounted on a 1.5 mm clean coverglass (EMS # 72200-31) precoated with 1 mg/ml BSA:BSA-Biotin (9:1 molar ratio) for 20 min at room temperature, washed in a water bath and then overnight at room temperature with 5 μg/ml Neutravidin in PBS. Extra Neutravidin was also washed with water. The set up was then assembled with an Attofluor chamber. GUVs were immobilized for 10 min on BSA:BSA-Biotin and Neutravidin coated clean cover glasses. 7.5 μM mVP40 proteins or His-tag alone were incubated with 50 μg/ml Ni-NTA-Atto 550 dye (Millipore Sigma, Burlington, MA) in a final volume of 500 μl, overnight at 4°C. Prior to incubation with GUVs, the proteins were concentrated to 100 μl using 30K MWCO concentration tubes (or 3K MWCO for His-tag alone purification). This step allowed to remove extra Ni-NTA-Atto 550 not bound to the proteins. The GUVs and proteins are then incubated for 30 min at 37°C at protein final protein concentration of 1.5 μM with the GUVs. N&B analysis was performed with the similar set up described above with some optimization. Briefly, at least 100 frames were imaged with Zeiss LSM 880 upright microscope using a Plan Apochromat 63x 1.4 numerical aperture oil objective, laser power: 0.1% using 561 nm laser, image size 256 × 256 pixel, pinhole: 4 μm, scan speed: 8.19 drop μsec, 16 bit depth.

On each experimental day, free NTA-Atto550 dye with GUVs was imaged and the true brightness (B) of a monomeric dye was determined (0.075-0.098). The apparent brightness value of mVP40 oligomers was calculate as described above using SimFCS software. Bins were placed in the brightness plot to correspond with the respective oligomer size. The number of pixels of monomer-dimer, dimer-hexamer, hexamer-12mer, 12mer-24mer, and 24mer+ bins were recorded. Average % pixels of each oligomeric state at the GUV membrane was ratiometrically determined by the total number of pixels in each bin vs. the total number of pixels in the image

### MLV sedimentation assay and in Vitro crosslinking reaction

MLV sedimentation assays were performed as described previously (Wijesinghe and Stahelin, 2016). For *in vitro* crosslinking assays, 0.2 mM LUVs were used. 2 μM of mVP40 wild type or mutant proteins were allowed to incubate with LUVs of four different lipid compositions (100% PC, PC:PS (70:30), PC:PIP_2_ (92.5:7.5), PC:PS:PIP_2_ (75:20:5) for 20 minutes at room temperature with a total reaction volume of 50 μl. Crosslinking agent – BS^3^ (ThermoFisher Scientific) was added to a final concentration of 1 mM and allowed to incubate with the lipid-protein mixture for 30 minutes. Reactions were stopped by adding 1 μl of glycine to a final concentration of 50 mM for 15 minutes at room temperature. 20 μl from each reaction was run on a SDS-PAGE gel and the protein bands were observed using silver staining.

### HDX-MS analysis

HDX-MS analysis of W83R/N148A, L226R/S229R and WNL mutants in the presence and absence of anionic lipid vesicles (PC:PS 55:45) was conducted as described in (Wijesinghe *et al.*, 2017).

### Molecular Dynamics Simulations

The mVP40 hexamer structure was modeled based on the eVP40 hexamer (PDB ID: 4LDD) as the template. The modeled mVP40 hexamer was relaxed with all atom molecular dynamics simulation using NAMD2.12. For this, an mVP40 hexamer system was set up using *Charmm gui* solvation builder^4^. The system was solvated using TIP3 water molecules in 0.15 M KCl. The simulation was performed with Charmm36m force fields^5^ and a SHAKE algorithm was used to treat covalent atoms whereas pressure was maintained using the Nose-Hoover Langevin-piston method. Similarly, the particle mesh Ewald (PME) method was used for the long-range electrostatic interactions. After 10,000 steps of minimization and 200 ps equilibration, production simulation was performed for 100 ns at 300 K using 2 fs time step. Additionally, two NTDs making up an oligomer interface was simulated for 150 ns. VMD was used to analyze the trajectories and protein images.

## General

S.A., M.L.H., and R.V.S. thank Nathan J. Dissinger for excellent technical support

## Author contributions

S.A., M.L.H. and K.J.W. designed and performed research; S.M.A., B.S.G, P.P.C., N.B. and S.L. performed research and wrote corresponding sections; all authors analyzed data. S.A, M.L.H and R.V.S wrote the paper and all the authors agreed on the submitted manuscript.

## Supplementary Materials

**Table S1.**
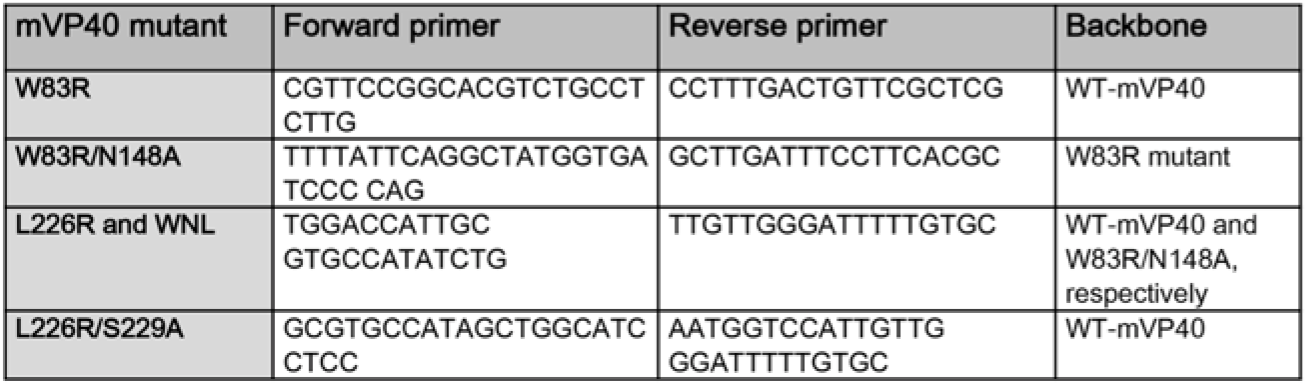
List of primers used to generate mVP40 mutant using site directed mutagenesis.

**Table S2.**
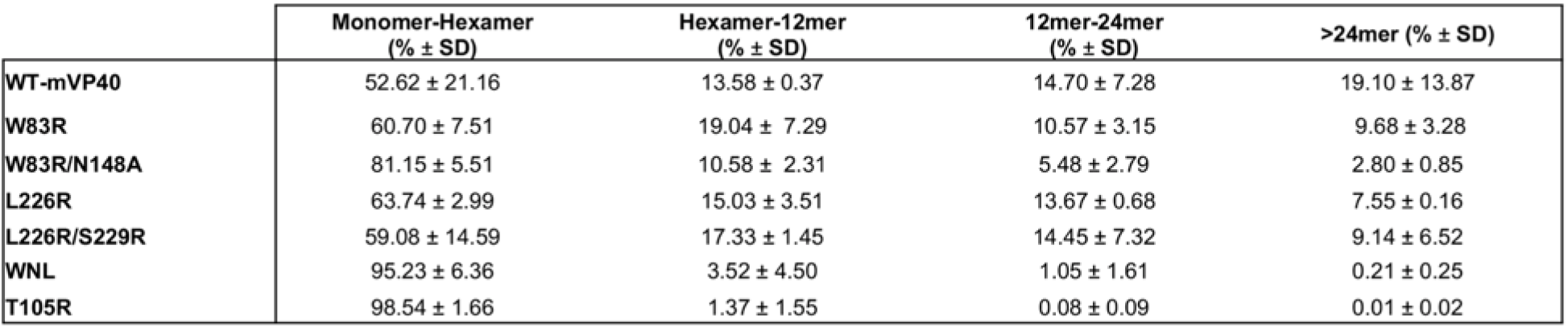
Summary of live cell Number and Brightness analysis on eGFP-mVP40 expressing cells.

|  | Monomer-Hexamer<br>(% ± SD) | Hexamer-12mer<br>(% ± SD) | 12mer-24mer<br>(% ± SD) | >24mer (% ± SD) |
| --- | --- | --- | --- | --- |
| WT-mVP40 | 52.62 ± 21.16 | 13.58 ± 0.37 | 14.70 ± 7.28 | 19.10 ± 13.87 |
| W83R | 60.70 ± 7.51 | 19.04 ± 7.29 | 10.57 ± 3.15 | 9.68 ± 3.28 |
| W83R/N148A | 81.15 ± 5.51 | 10.58 ± 2.31 | 5.48 ± 2.79 | 2.80 ± 0.85 |
| L226R | 63.74 ± 2.99 | 15.03 ± 3.51 | 13.67 ± 0.68 | 7.55 ± 0.16 |
| L226R/S229R | 59.08 ± 14.59 | 17.33 ± 1.45 | 14.45 ± 7.32 | 9.14 ± 6.52 |
| WNL | 95.23 ± 6.36 | 3.52 ± 4.50 | 1.05 ± 1.61 | 0.21 ± 0.25 |
| T105R | 98.54 ± 1.66 | 1.37 ± 1.55 | 0.08 ± 0.09 | 0.01 ± 0.02 |

**Table S3.** Summary of Number and Brightness analysis on GUVs incubated with Ni-NTA-Atto550 conjugated 6cHis-mVP40 proteins.

|  | % bound protein ± SD |  |  | Monomer-Dimer (mean % over total bound protein) |  |  | Dimer-Hexamer (mean % over total bound protein) |  |  | Hexamer-12mer (mean % over total bound protein) |  |  | 12mer-24mer (mean % over total bound protein) |  |  | >24mer (mean % over total bound protein) |  |  |
| --- | --- | --- | --- | --- | --- | --- | --- | --- | --- | --- | --- | --- | --- | --- | --- | --- | --- | --- |
|  | PS | PI(4,5)P<sub>2</sub> | PS:PI(4,5)P<sub>2</sub> | PS | PI(4,5)P<sub>2</sub> | PS:PI(4,5)P<sub>2</sub> | PS | PI(4,5)P<sub>2</sub> | PS:PI(4,5)P<sub>2</sub> | PS | PI(4,5)P<sub>2</sub> | PS:PI(4,5)P<sub>2</sub> | PS | PI(4,5)P<sub>2</sub> | PS:PI(4,5)P<sub>2</sub> | PS | PI(4,5)P<sub>2</sub> | PS:PI(4,5)P<sub>2</sub> |
| WT-mVP40 | 14.57 ± 11.85 | 8.89 ± 7.8 | 22.43 ± 19.2 | 53.60 | 56.54 | 17.76 | 36.93 | 24.72 | 30.29 | 9.40 | 14.25 | 31.44 | 0.07 | 4.25 | 16.08 | 0.00 | 0.23 | 4.43 |
| W83R/N148A | 20.38 ± 12.13 | 11.73 ± 13.61 | 6.62 ± 8.17 | 79.49 | 50.83 | 13.31 | 18.78 | 26.36 | 30.51 | 1.73 | 16.27 | 33.31 | 0.00 | 5.60 | 20.70 | 0.00 | 0.92 | 2.17 |
| L226R | 20.63 ± 10.91 | 21.81 ± 12.02 | 14.02 ± 18.4 | 69.07 | 45.49 | 13.15 | 25.11 | 28.87 | 51.36 | 5.72 | 17.49 | 23.77 | 0.10 | 6.04 | 8.37 | 0.00 | 2.12 | 3.35 |
| WNL | 21.65 ± 15.18 | 13.78 ± 12.29 | 13.34 ± 12.53 | 78.52 | 57.70 | 11.36 | 18.40 | 27.66 | 30.88 | 2.56 | 11.57 | 37.78 | 0.52 | 2.39 | 17.40 | 0.00 | 0.69 | 2.58 |
| T105R | 0.02 ± 0.07 | 1.03 ± 1.36 | 0.55 ± 1.64 | - | - | - | - | - | - | - | - | - | - | - | - | - | - | - |

**Figure S1.**
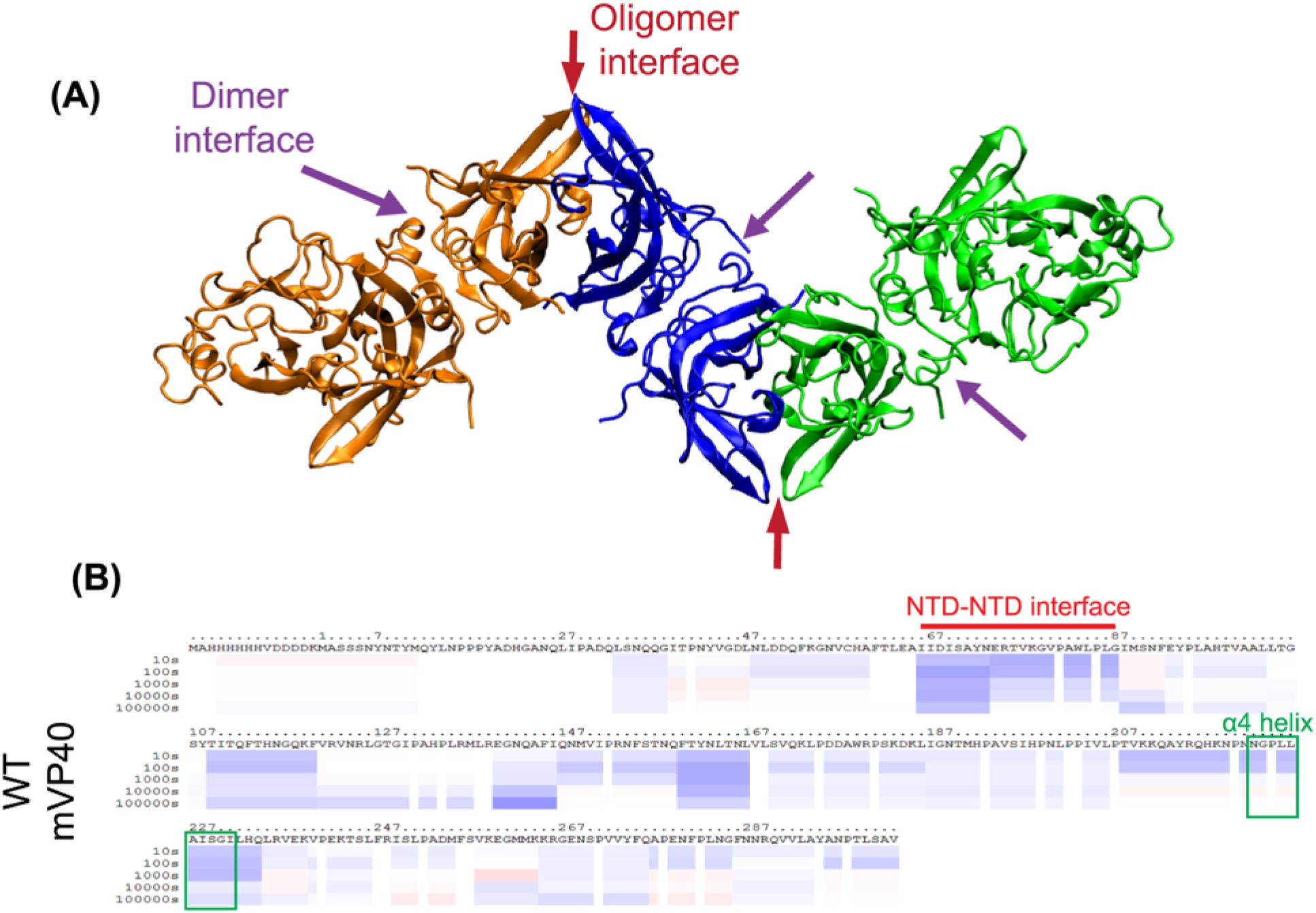
mVP40 potential oligomerization through binding to PS containing lipid vesicles. Structure of the mVP40 hexamer showing different modeled interfaces based on the Ebola VP40 (eVP40) hexamer (PDB ID: 4LDD). Each dimer is colored differently. (B) Differences in deuteration level (D%) in the presence of PS-containing liposomes mapped to mVP40 protein sequence. Each row corresponds to the exchange level compared to mVP40 in the absence of liposomes from 10 to 100,000 s. Color coding: blue indicates the regions that exchange slower in the presence of lipid and red adapted from *a* research originally published in the Journal of Biological Chemistry. Kaveesha J. Wijesinghe, Sarah Urata, Nisha Bhattarai, Edgar E. Kooijman, Bernard S. Gerstman, Prem P. Chapagain, Sheng Li, and Robert V. Stahelin. Detection of lipid-induced structural changes of the Marburg virus matrix protein VP40 using hydrogen/deuterium exchange-mass spectrometry. J Biol Chem. 2017; 292:6108-6122. © the American Society for Biochemistry and Molecular Biology (Wijesinghe *et al.*, 2017).

**Figure S2.**
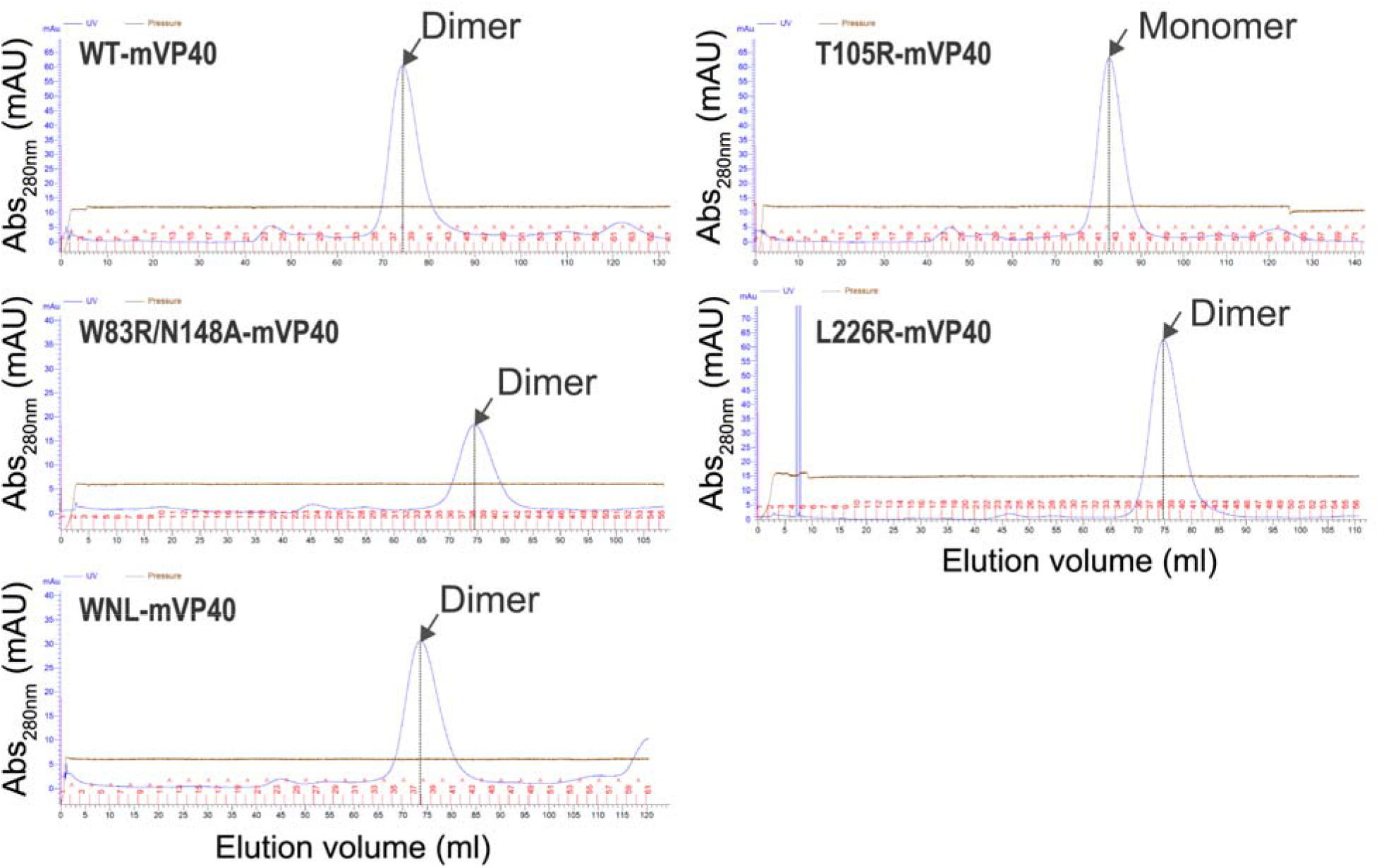
Gel filtration analysis of mVP40 oligomerization interfaces. Size exclusion chromatographs (SEC) of mVP40 wild type (WT), W83R/N148A, WNL and L226R mutants shown as absorbance (280nm) versus elution volume. In brief, proteins post Ni-NTA purification were injected onto HiLoad® 16/600 Superdex® 200 pg column. Dimeric mVP40 is eluted at elution volume from 68 ml to 82 ml. T105R-mVP40 is eluted as a monomer at elution volumes from 75 ml to 90 ml.

**Figure S3.**
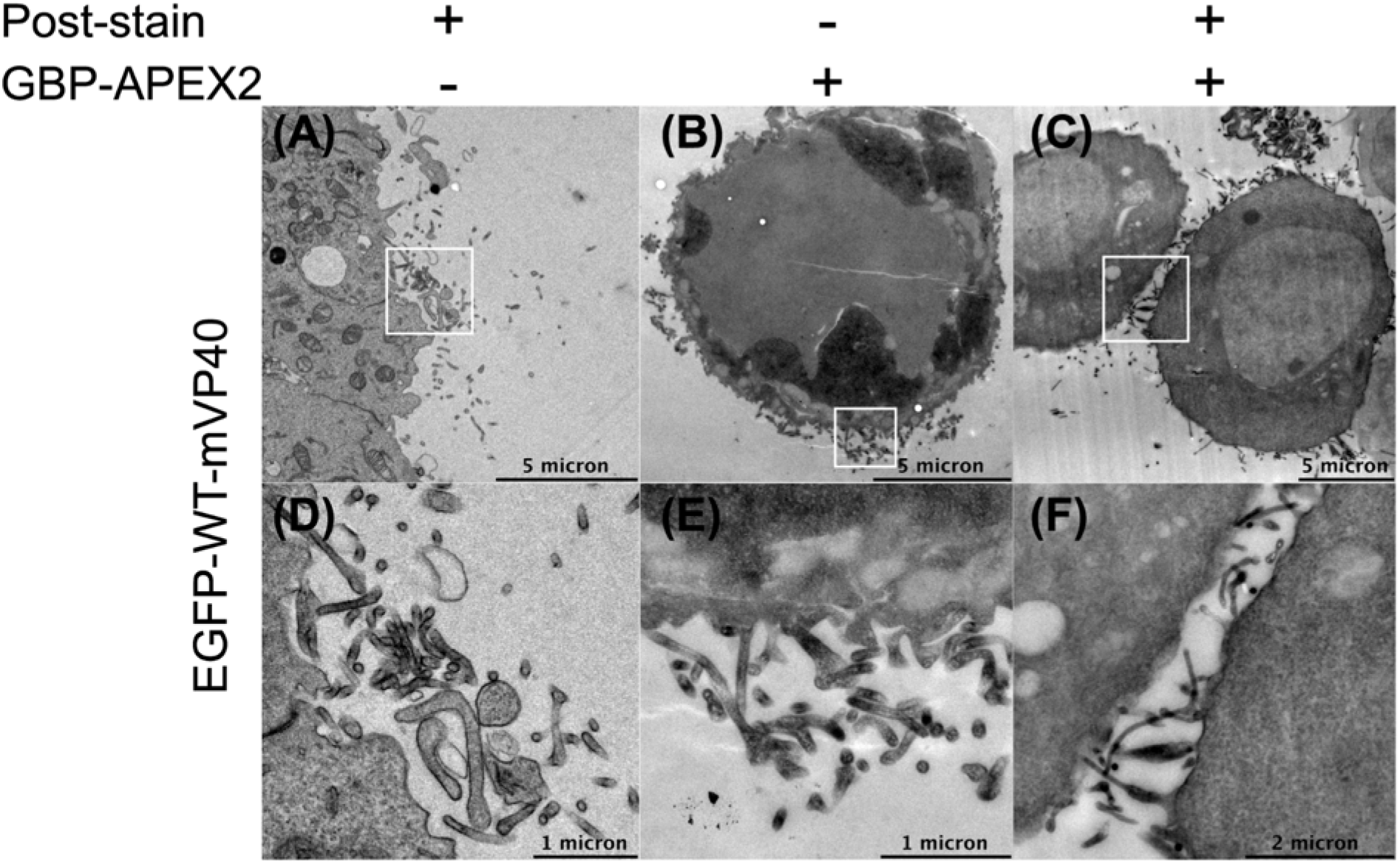
EGFP-mVP40 displays typical membrane elongation and budding of filamentous VLPs at the surface of HEK293 cells co-expressing GBP-APEX2. Electron microscopy micrographs of cells expressing different plasmids: **(A)** EGFP-mVP40 alone or **(B)**, **(C)** with GBP-APEX2, for 14 hours before chemical fixation, post-stained **(A)** and **(C)** or not **(B)** prior to imaging. **(D)**, **(E)** and **(F)** are zoomed insets from **(A)**, **(B)** and **(C)**, respectively.

**Figure S4.**
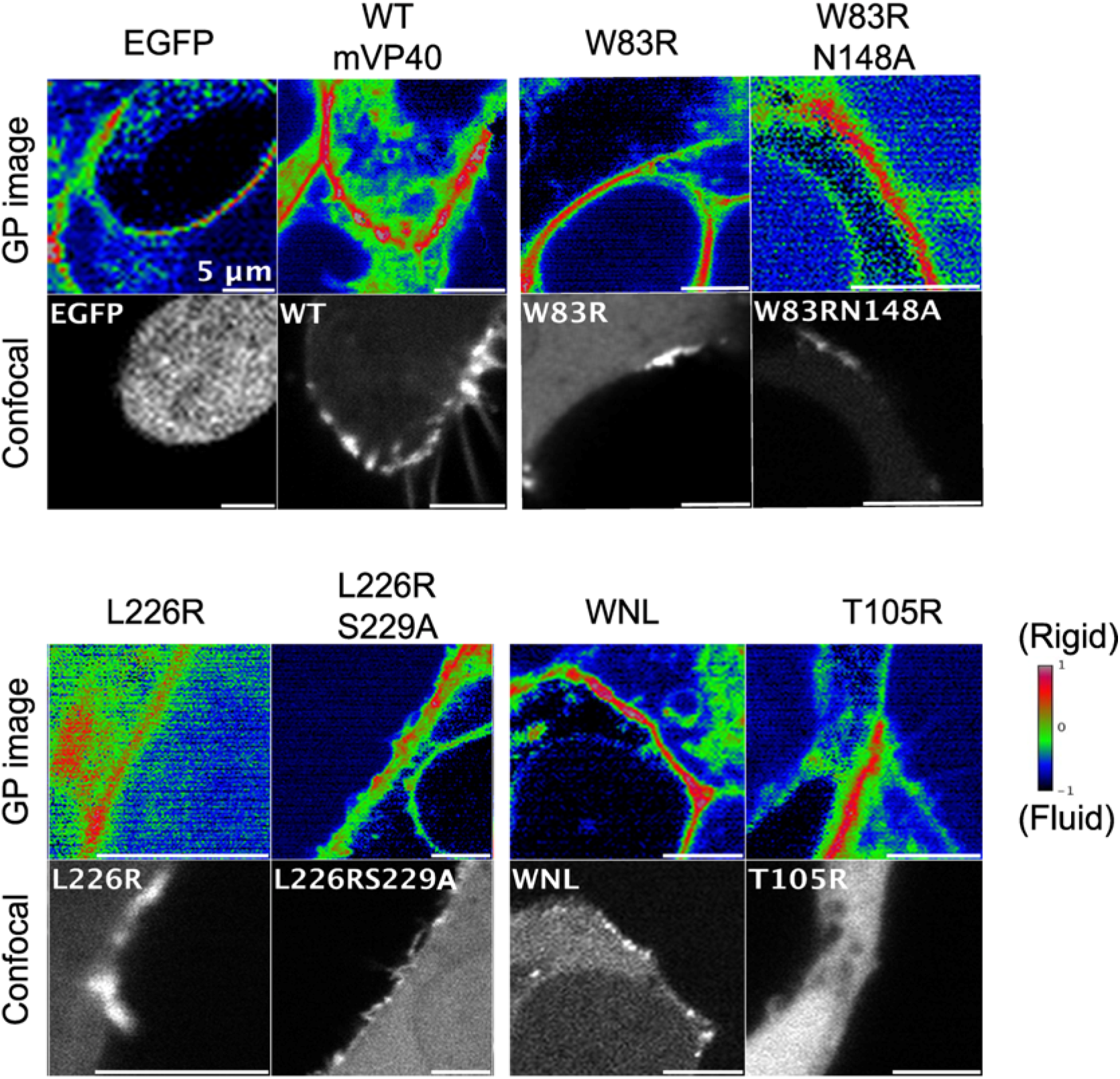
Laurdan general polarization (GP) images of HEK293 cells expressing EGFP-mVP40. Relationship between GP value and EGFP signal distributions across the plasma membrane. HEK293 cells were incubated with 10 μM laurdan dye 14 h.p.t with EGFP constructs. Multiphoton (top panel) and confocal imaging (bottom panel) were performed after 30 min incubation with the dye. Color coding: red indicates rigid membrane while blue indicates fluid membranes.

**Figure S5.**
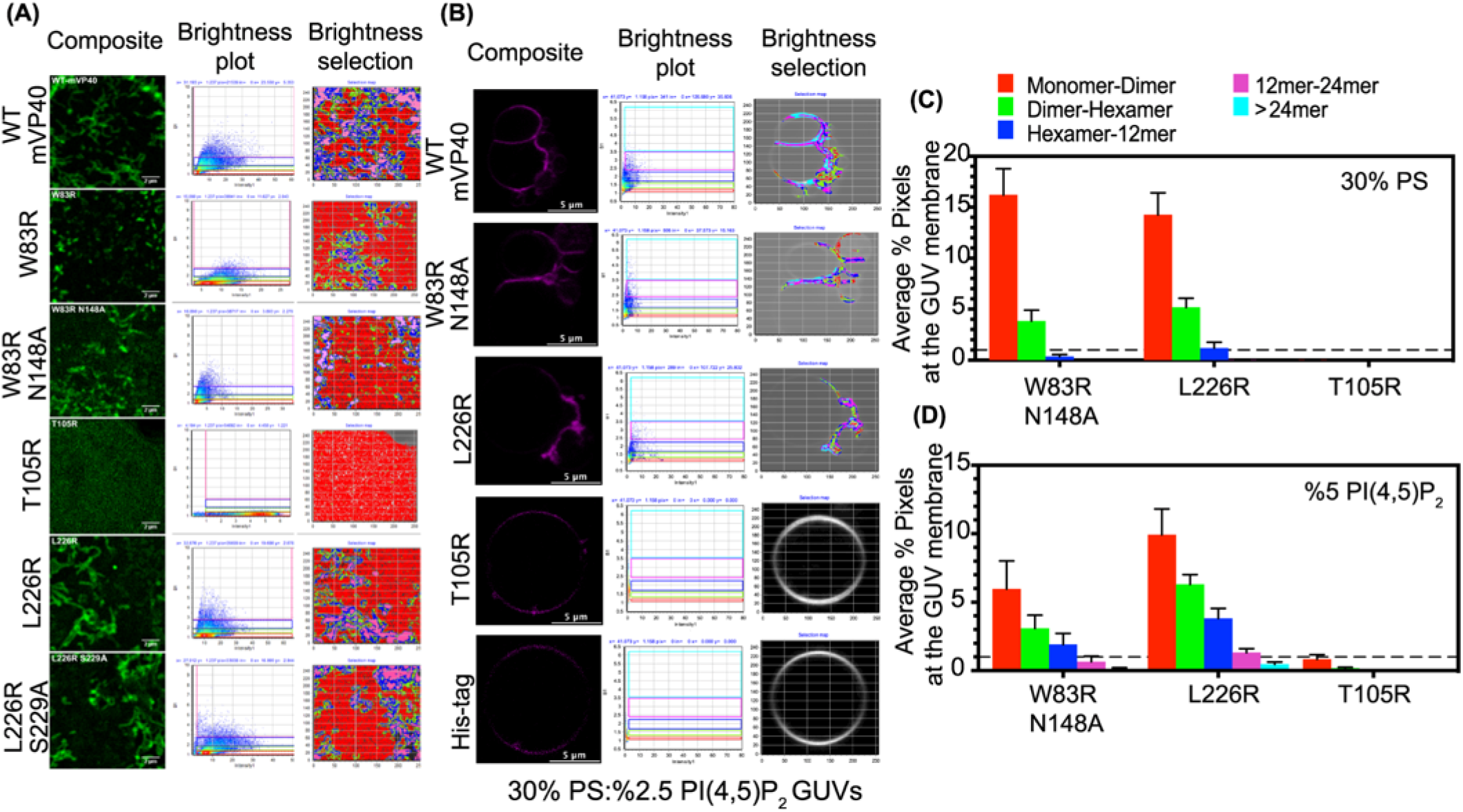
Cellular and *in vitro* oligomerization profiles of mVP40 mutants analyzed through Number & Brightness analysis. HEK293 cells transiently expressing GFP-fused mVP40 constructs **(A)** or GUV containing 30%PS:2.5% PI(4,5)P_2_ incubated with 6xHis tagged protein coupled to Ni-NTA-Atto 550 **(B)** were imaged and Number & Brightness (N&B) analysis was performed using SimFCS software. Representative images of the workflow in SimFCS for N&B analysis of EGFP-WT-mVP40, EGFP-W83R-mVP40, EGFP-W83R/N148A-mVP40, EGFP-T105R-mVP40, EGFP-L226R-mVP40, EGFP-L226R/S229R-mVP40 are shown in (A) and in (B) from N&B analysis of GUV 6xHis-WT-mVP40, 6xHis-W83R/N148A-mVP40, 6xHis-T105R-mVP40, 6xHis-L226R-mVP40 and 6xHis tag alone. The original composite of the time-lapse images (left panel), the number of pixels vs. intensity plot (middle panel) and brightness selection plot of the cell (right panel) are shown for each analysis. **(C)** Oligomerization profiles of W83R/N148A, L226R and T105R mutants according to different anionic membranes 30% PS only and 5% PI(4,5)P_2_ only, determined from N&B analysis. Values are reported as mean ± S.E.M of three independent means.

**Figure S6.**
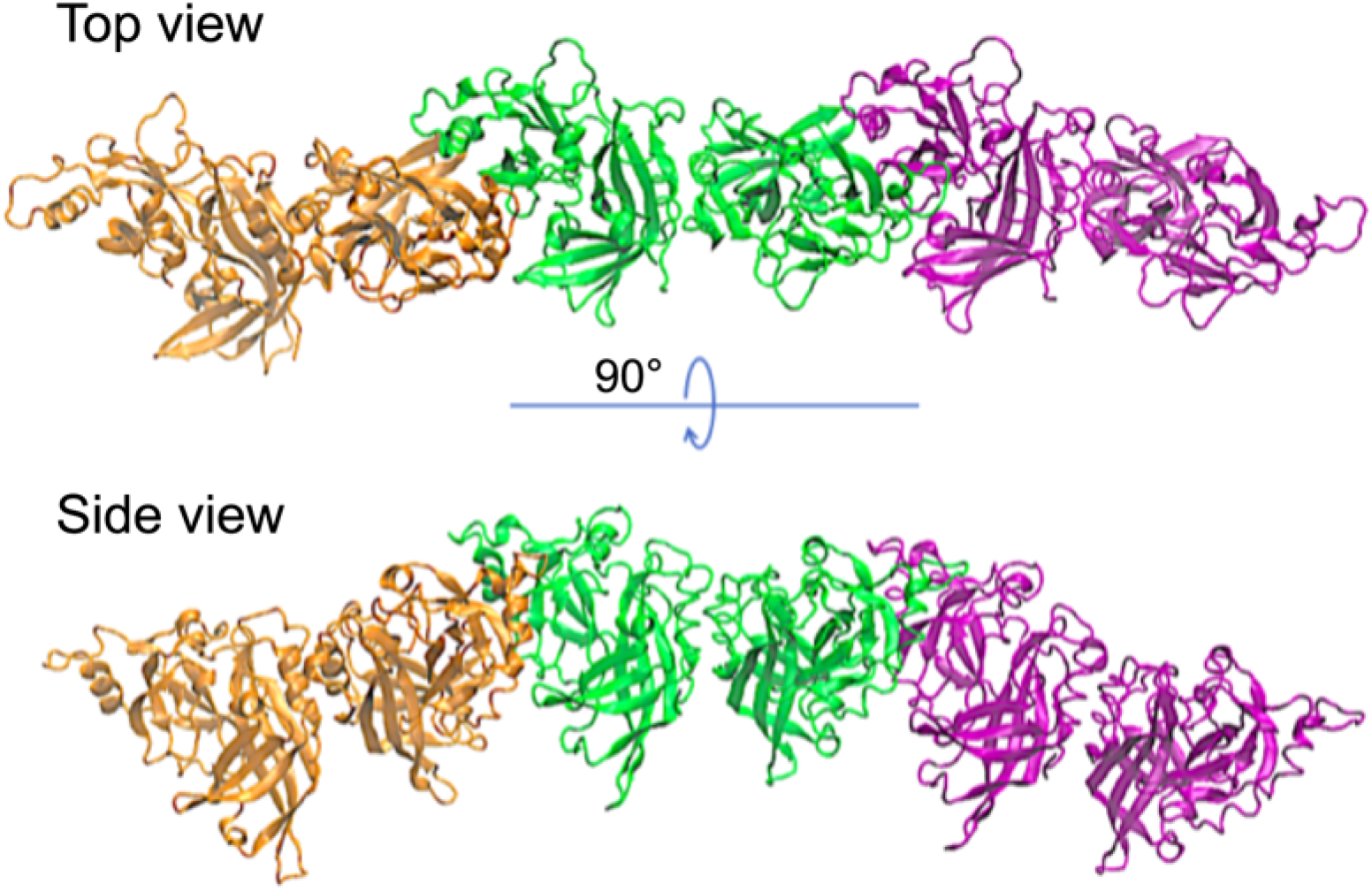
MD simulation of mVP40 potential CTD-CTD linear oligomerization. Each mVP40 dimer is represented by a single color.

**Movie S1. Molecular Dynamics Simulation of NTD oligomerization interfaces in mVP40 compared to eVP40.** The oligomerization of mVP40 through interactions of W83 residues at NTD is mediated by relaxation of NTD region of each protein. In contrast, the distance between W95 residues in eVP40 does not change over time during the MD simulation.

